# RiboCop surveils pre-rRNA processing by Dicer in cellular quiescence

**DOI:** 10.1101/2025.09.16.676644

**Authors:** B. Roche, R.A. Martienssen

## Abstract

In nature, most cells exist in a quiescent G_0_ state in which cellular homeostasis must be rigorously maintained in the absence of cell division. Non-coding RNAs are prevalent in G_0_ and are important regulators of development and differentiation, but their function in quiescence is unclear. Here, we identify pre-rRNA as a direct target of the RNase III enzyme Dicer specifically in quiescence. Dicer is physically present at the rDNA, and improper rRNA processing in mutants results in a nucleolar stress response involving a novel *trans*-acting non-coding RNA (RiboCop) in complex with the highly conserved proteins Enp2/NOL10 and RNase H1. RiboCop is complementary to unprocessed pre-rRNA and triggers rDNA repeat silencing via Sir2, RENT, and histone H3-lysine-9 (H3K9) methylation. Thus RiboCop silences rDNA specifically during dormancy, when silencing of non-functional rRNA becomes essential.

## Main Text

Cellular quiescence is the state of non-dividing cells that are still metabolically active and able to re-enter the cell cycle when given the appropriate signal; this state is common in nature, ranging from unicellular yeasts to most stem cells in mammals *(1–4)*. Despite the importance and evolutionary conservation of quiescence, relatively little is known about the molecular mechanisms underlying its establishment and maintenance. A common feature is strong down-regulation of overall transcription to basal levels while still expressing a wide diversity of transcripts *(5,6)*. In multicellular organisms, the stem cell niche is an additional major contributor to the maintenance of quiescence via extracellular signals *(7,8)*. This complicates our understanding of intracellular quiescence pathways, which has largely advanced by studying unicellular model organisms such as yeasts. In particular, the fission yeast *Schizosaccharomyces pombe* is a very well-suited model organism to study cellular quiescence, because it can be triggered by a simple signal—nitrogen-starvation of a prototrophic strain—in a near-homogenous and synchronous manner *(9,10)*.

While expressed at low levels, non-coding RNAs (ncRNAs) occupy a much larger proportion of the transcriptomic diversity in *S. pombe* G_0_ cells *(11)*, raising the possibility that they contribute significantly to the transcriptional reprogramming of quiescence. This prevalence of ncRNAs in quiescence appears to be a conserved feature in evolution, also observed from budding yeast *(12)* to dormant cancer cells *(13)*. Specific ncRNAs may be essential to control gene expression to adapt to the quiescent transcriptome; yet, most individual ncRNA mutants do not display phenotypes in quiescent cells *(14)*, indicating significant redundancy and suggesting the possibility that it is their global regulation that is required, particularly by RNA surveillance pathways. In accordance with this idea, we have previously found that RNA interference (RNAi) becomes essential specifically during quiescence in *S. pombe (15)*, and recent studies have also highlighted the essential role of RNA surveillance pathways *(11,16–18)*.

In the absence of RNAi, cells quickly lose viability during quiescence maintenance, due to the accumulation of stalled RNA pol I and H3K9me2 at the repetitive rDNA locus *(15)*. However, the nature of the RNAi target causing this phenotype has been unclear. In this study, we identify the pre-ribosomal RNA (pre-rRNA) as the Dicer target in quiescent cells, and we find that Dicer directly binds to rDNA chromatin. Improper pre-rRNA processing in Dicer mutants results in RNA pol I elongation defects and a nucleolar stress response. We further identify the novel ncRNA RiboCop as the mediator recruiting silencing factors to rDNA during this nucleolar stress response. Unlike the non-coding *cis*-acting pRNA (promoter RNA) in mammals *(20)*, RiboCop acts in *trans* to silence rDNA by recognizing unprocessed pre-rRNA and recruiting rDNA silencing factors, in accordance with a global role for RNA surveillance pathways in maintaining quiescence and silencing rDNA in dormant cells.

### Detection of Dicer cleavage sites in quiescence by iPARE-seq

We have previously found that RNAi becomes essential in *S. pombe* specifically in quiescence and that RNAi mutants, such as Dicer (*dcr1*Δ) and Argonaute (*ago1*Δ), accumulate lethal amounts of heterochromatin at the repetitive rDNA locus *(15)*. Because the catalytic mutant *dcr1-5* (D908A, D1127A) *(21)* displays the same G_0_-defective phenotype as *dcr1*Δ (Fig. 1A), we reasoned that Dicer must exert its function *via* cleavage of RNA substrates. We could not detect novel Dicer-dependent small RNAs in quiescent cells *(15)* (fig. S1), suggesting the target could be the rRNA itself. We therefore aimed to identify candidate RNA targets of Dicer in G_0_ in a direct manner by sequencing the degradome of wild-type and *dcr1*Δ G_0_ cells. We designed iPARE-seq, an improved degradome-sequencing technique derived from PARE-seq (Parallel Sequencing of RNA ends) *(22)*, in which available 5’-phosphate ends in the transcriptome are ligated to a biotinylated adapter followed by purification, library generation and sequencing (fig. S1B). Endonucleolytic cleavage by RNase III-family enzymes results in a 5’P end, as shown for *E. coli* RNase III, *S. pombe* Rnt1(=Pac1) and Dicer enzymes *(23–25)*, leading us to expect that cleaved Dicer targets would be recovered using this approach. In addition to 5’P-containing cleaved RNA pol II transcripts, RNA pol I and RNA pol III transcripts are also recovered as they do not harbor a 5’ 7-methylguanidine cap, along with their major processed forms (fig. S1C). iPARE-seq was performed in wild-type and *dcr1*Δ G_0_ cells (n=3), as well as in the catalytic-dead mutant *dcr1-5* (n=2). As expected, the strongest 5’P peaks are observed for mature rRNA ends, which correspond to the known A_1_, B_1L/S_ and C_1_ cleavage sites (determining the 5’ end of mature 18S, 5.8S and 28S rRNA respectively *(26)*) (Fig. 1B), and we could recover these cleavage sites by iPARE-seq with single-nucleotide precision (fig. S1D). In *dcr1*Δ cells, these peaks are greatly reduced, consistently with a ribosomal RNA accumulation defect (Fig. 1B; fig. S1F).

**Fig. 1.**
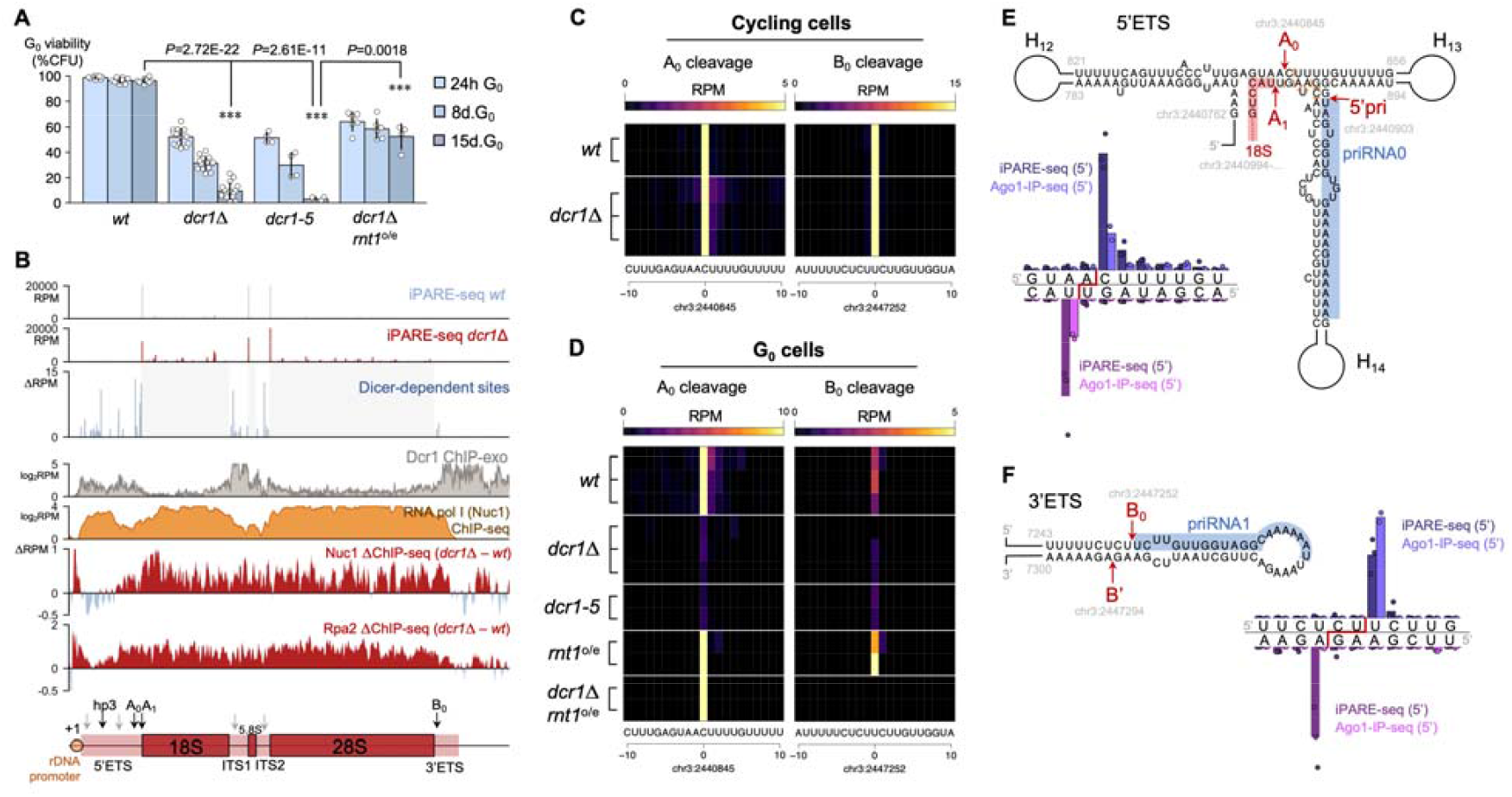
Dicer and Rnt1 process pre-ribosomal RNA in quiescent cells. (**A**) The *dcr1*Δ deletion mutant and *dcr1-5* catalytic mutant lose viability during quiescence. Overexpression of Rnt1 in G_0_ (*p*.*urg1* promoter) results in phenotypic suppression and maintained viability over 15 days of quiescence (*** represents *P*<0.01, t-test). (**B**) Mapping of accessible 5’-monophosphate ends by iPARE-seq in wild-type and *dcr1*Δ quiescent cells uncovers a set of Dicer-dependent cleavage events over the pre-rRNA spacer elements. Dcr1 binds the rDNA, in particular the spacers, as seen by ChIP-exo (*dcr1*-TwinStrepII background). Nuc1^A190^ (RNA pol I) ChIP-seq (*nuc1*-(Gly)_6_-3xFLAG background) provided for comparison. Differential ChIP-seq shows an accumulation of RNA pol I (Nuc1^A190^, Rpa2^A135^) in *dcr1*Δ mutants both at the rDNA promoter, as well as over the entire rDNA repeat, starting within the last third of the 5’ETS region. (**C**) iPARE-seq recovers the key A_0_ and B_0_ pre-rRNA cleavage sites with single-nucleotide precision, and Dcr1 is dispensable for either cleavage in cycling cells; whereas (**D**) in quiescent cells, both A_0_ and B_0_ become Dcr1-dependent. Overexpression of Rnt1 rescues A_0_ cleavage (but not B_0_) in *dcr1*Δ mutants. (**E**) The 5’ETS folds into extended dsRNA hairpins, showing the co-localization of A_0_ and A_1_ cleavage sites, in proximity to Argonaute-bound priRNA0. Both iPARE-seq and 5’-end coverage in G_0_ Ago1-RNA-IP-seq recover A_0_ and A_1_ sites with single-nucleotide precision. (**F**) The 3’ETS folds into a hairpin with B_0_ and B’ cleavage sites. iPARE-seq and Ago1-IP-seq (5’) recover these sites with single-nucleotide precision. Cleavages with a 3’-overhand 1-2nt offset are compatible with processing by RNase III family endonucleases.

Many iPARE-seq sites were found within the rDNA locus, with a non-random distribution and high correlation between replicates (Fig. 1B). Peaks in the 18S, 5.8S and 28S sequences are mostly due to degradation of mature rRNA, and accumulate in *dcr1*Δ cells in accordance with an RNA pol I stalling defect and a loss of viability during quiescence *(15)*. We therefore focused on pre-rRNA cleavage sites, which are absent from mature rRNA molecules, and which constitute key steps in the regulation of eukaryotic pre-rRNA processing and ribosome biogenesis *(26)*. We recovered two early cleavage sites, A_0_ and B_0_ in the 5’ and 3’ External Transcribed Spacers (ETS) respectively, as well as the +1 Pol I-transcriptional start site (TSS), with single-nucleotide precision (Fig. 1BCD, fig. S1D). In addition, we identified 13 new pre-rRNA cleavage sites in the 5’ETS. Cleavage at all but one of these sites was strongly down-regulated in *dcr1*Δ cells, indicating that pre-rRNA processing is defective in *dcr1*Δ (Fig. 1B, fig. S1F). The most striking loss was at the key A_0_ cleavage site, which is reduced to 10% in *dcr1*Δ cells (t-test *P*=0.0079) and 9% in the catalytic mutant *dcr1-5* (t-test *P*=0.0075). In contrast, the iPARE-seq cleavage profile of cycling cells indicated that Dicer is dispensable for A_0_ cleavage (Fig. 1CD). In the 3’ETS, the B_0_ cleavage site, which is cleaved in cycling cells by Rnt1 *(27)* is also reduced (21% in *dcr1*Δ) indicating that cleavage at B_0_ also becomes Dicer-dependent specifically in G_0_ cells (Fig. 1D).

### RNase III switches between Dicer and Rnt1 between proliferation and quiescence

A_0_ is the main 5’ETS cleavage site, directed by the U3 snoRNA and its associated ribonucleoprotein complex *(28)*, conserved from yeast to mammals *(26,29)*. In *S. cerevisiae*, which lacks Dicer, RNT1 has been proposed as the A_0_ nuclease, as it displayed the ability to cleave a minimal *in vitro* A_0_ substrate *(30)*, but may not be sufficient as residual cleavage is detected in viable *rnt1*Δ mutants *(31)*. In *S. pombe, rnt1*^*+*^ is essential, precluding us from analyzing *rnt1*Δ deletion mutants for their contribution to G_0_ pre-rRNA processing. We opted for an alternative strategy: overexpressing *Rnt1*^*+*^ in G_0_ cells, using the *p*.*urg1*_800_ promoter, which is activated not only by uracil but also during quiescence entry *(32)*, thus allowing G_0_ overexpression *(15)*. The resulting *p*.*urg1::rnt1* strain (*rnt1*-o/e) is fully viable in quiescence.

Strikingly, we found that the *dcr1*Δ *purg1::rnt1* strain strongly rescued the G_0_ maintenance defects normally seen in *dcr1*Δ (Fig. 1A), similarly to class II suppressors such as *dcr1*Δ*clr4*Δ *(15)*. This suggests that Rnt1^+^ can compensate for Dcr1^+^ in G_0_ cells when expressed at sufficient levels. To ascertain whether this suppression is caused by rescue of pre-rRNA processing, we performed iPARE-seq in *purg1::rnt1* and in *dcr1*Δ *purg1::rnt1* strains, and found that several Dicer-dependent pre-rRNA cleavage sites were indeed restored in these strains, including A_0_ (Fig. 1D). Moreover, the main Rnt1 target B_0_ was also processed at higher levels in *purg1::rnt1* as expected (Fig. 1D). Recent models have proposed that A_0_ is cleaved in a co-transcriptional manner, while B_0_ is processed after the full-length pre-rRNA is transcribed, and that the balance between these processing sites is dependent on RNA pol I elongation speed, growth phase and nutrient availability *(26,33)*. In accordance with these models, B_0_ cleavage is prevalent in cycling cells, but shifts to A_0_ processing in quiescent cells (∼3.8-fold B_0_ preference in cycling cells *vs*. ∼6.8-fold preference for A_0_ in quiescent cells). We reasoned that the A_0_/B_0_ switch may reflect a specialization of Dcr1 vs Rnt1, with a differential requirement for pre-rRNA processing co-transcriptionally *vs*. post-transcriptionally.

RNase III-family enzymes cleave dsRNA, and so we hypothesized that A_0_ and B_0_ sites are locally folded in dsRNA structures (Fig. 1EF). While the exact folding and structure of the A_0_ site is unresolved in *S. cerevisiae* processome cryo-EM structures *(34)*, it can fold with neighboring hairpins to result in long dsRNA regions, a known substrate for RNase III enzymes like Dicer and Rnt1. A cruciform-like structure was proposed for *S. pombe*, and the presence of long dsRNA hairpins (H_12_ and H_13_) is essential for processing to occur on pre-rRNA plasmid templates *(35)*. Similar substrates with contiguous dsRNA regions were shown to be efficient substrates for human Dicer *(36)*. In this structure, the A_0_ and A_1_ sites occur in dsRNA. Both our G_0_ iPARE-seq and published G_0_ Ago-IP small RNA-seq *(17)* datasets provide evidence for 5’-monophosphate ends at A_0_ and A_1_, with a 3’-overhang 1-nt offset (Fig. 1E).

Re-analysis of the G_0_ Ago-IP small RNA-seq dataset identified priRNA1, a small RNA derived from pre-rRNA which is loaded into Ago1 in both cycling cells *(37)* and quiescent cells *(17)*, as expected, but also a second highly-prevalent Ago1-loaded priRNA (which we termed priRNA0) located immediately proximal to the A_0_/A_1_ site, suggesting the physical presence of not only Dcr1 but also Ago1. Correct processing may require further dsRNA formation, as we observed that A_0_ cleavage is also inhibited in the catalytic-dead RNA-dependent RNA polymerase mutant *rdp1*-D903A (fig. S1F), confirming the requirement of all three RNAi factors (Dcr1, Ago1, Rdp1) for pre-rRNA processing in quiescent cells. Likewise, the B_0_ cleavage site is located on a dsRNA hairpin, with an associated cleavage (B’) on the antisense strand and a canonical 3’-overhang 2-nt offset (Fig. 1F), matching previously identified Rnt1 3’ETS cleavage in cycling cells *(27)*. The sense strand corresponds exactly to the 5’ end of priRNA1. Consistent with B_0_ cleavage by both Dcr1 and Rnt1, priRNA1 is Dicer-independent *(37)*. Overall, these results suggest that pre-rRNA is a major RNA target of Dicer specifically in G_0_.

### Dicer binds ribosomal RNA and prevents stalling of RNA pol I

Next, we aimed to determine the consequences of the pre-rRNA processing defect on rRNA transcription. We previously showed that in quiescence, *dcr1*Δ mutants display an increased occupancy of RNA pol I at the rDNA promoter *(15)*. To analyze the pattern of RNA pol I stalling, we performed ChIP-seq of the main RNA pol I subunits Nuc1^A190^ and Rpa2^A135^. As expected, RNA pol I binds across the full-length of the transcribed rDNA region (Fig. 1B). We found that in addition to the promoter, the accumulation of RNA pol I in *dcr1*Δ mutants starts specifically over the last third of the 5’ETS sequence where the A_0_ site is located, and then subsequently covers the entire rDNA repeat (Fig 1B), matching the region where H3K9 methylation accumulates in *dcr1*Δ G_0_ cells *(15)*. This suggests that the RNA pol I defect in *dcr1*Δ G_0_ cells is a consequence of the pre-rRNA processing defect that stalls Pol I. By analogy, defects in Coilin/Mug174 (essential for Cajal body formation) also result in both accumulation of stalled RNA pol I and increased H3K9me *(18)*.

The requirement of Dcr1 for co-transcriptional processing of pre-rRNA strongly suggests its physical presence at rDNA chromatin. We performed ChIP-exo of C-terminally 3xFLAG-tagged Dicer, and found that both *dcr1*-3xFLAG and the catalytic-dead mutant *dcr1*-D908A, D1127A-3xFLAG were strongly associated with rDNA chromatin in G_0_ cells (fig. S2). We confirmed this result by performing ChIP-exo of N-terminally tagged Dicer with a twin-StrepII tag (twStrep-*dcr1* strain), which also strongly bound the rDNA repeat sequence (Fig. 1B; fig.

S2), Neither tagged strain affected the Dicer protein as they did not cause any G_0_ defect (fig. S2D). Interestingly, we did not detect *dcr1*-FLAG binding at centromeres, in accordance with previous observations *(38,39)*; however, we could detect binding of the catalytic-dead *dcr1-* D908A,D1127A-FLAG mutant, suggesting that Dcr1 is stabilized on its centromeric substrate in the absence of Dicing activity (i.e. frozen enzyme), in particular at the core centromeric region (comprising the *imr* repeats and the *cnt* region which binds CENP-A). Broader centromeric binding is seen as well in twStrep-*dcr1* (fig. S2), in which the N-terminal tag may affect Dicer’s helicase domain and stabilize its chromatin interaction. Overall, these results indicate that although Dicer’s role in cleaving centromeric ncRNAs is well-established, this process is likely only transiently associated with pericentromeric chromatin, as was suggested from previous Dcr1-DamID profiles *(40)*. By contrast, the binding pattern of Dicer at rDNA is strongly enriched in the transcribed spacer regions where Dcr1-dependent cleavage sites are found (Fig. 1B). Dicer binding to rDNA chromatin has also been observed in mouse embryonic stem cells, although its functional significance was not known *(41). S. pombe* Dcr1 contains a predicted C-terminal nucleolar localization signal, in accordance with its role at rDNA (fig. S1C).

### The long non-coding RNA NC30 mediates RNAi defects in G_0_

Forward genetic approaches, and in particular the selection of genetic suppressors, can provide important insights into molecular mechanisms. We used a microevolution strategy to obtain new spontaneous *dcr1*Δ G_0_ suppressors, as previously described *(15)*, reasoning that we could isolate upstream genes involved in nucleolar function and/or rRNA processing. In one new suppressor strain, we identified a large subterminal deletion (>139kb) of the right arm of chromosome 1, which we named *deltel1R*, encompassing 34 lncRNA and 46 protein-coding genes (Fig. 2AB; fig. S3AB). RNA-seq in *wt* and *dcr1*Δ G_0_ cells identified a limited number of differentially-expressed genes in *dcr1*Δ, mostly comprising upregulated lncRNAs (91/140; χ^2^-test, *P*=6E-33) (fig. S2DE), likely indirect targets as they did not generate Dicer-dependent small RNAs, unlike centromeric and subtelomeric transcripts (fig. S2FG). We focused on lncRNAs within the *deltel1R* interval to map the *deltel1R* suppressor, which we found to be one of the Dicer-dependent ncRNAs, NC30 (*SPNCRNA*.*30*) (Fig. 2C, fig. S3H; details on fine-mapping are provided in Methods). The double-mutant *dcr1*Δ*NC30*Δ suppressed *dcr1*Δ viability defects in G_0_ during quiescence maintenance, to the same extent as *dcr1*Δ*deltel1R* (Fig. 2B) and similarly to H3K9 methylation mutants such as *dcr1*Δ*clr4*Δ *(15)* or to the processing rescue mutant *dcr1*Δ*purg1::rnt1* (Fig. 1). Furthermore, both *deltel1R* and *NC30*Δ also suppressed the viability loss of *ago1*Δ and *rdp1*Δ in G_0_ (fig. S3J), indicating that NC30 is a general RNAi G_0_ suppressor (similarly to class ii suppressors *(15)*).

**Fig. 2.**
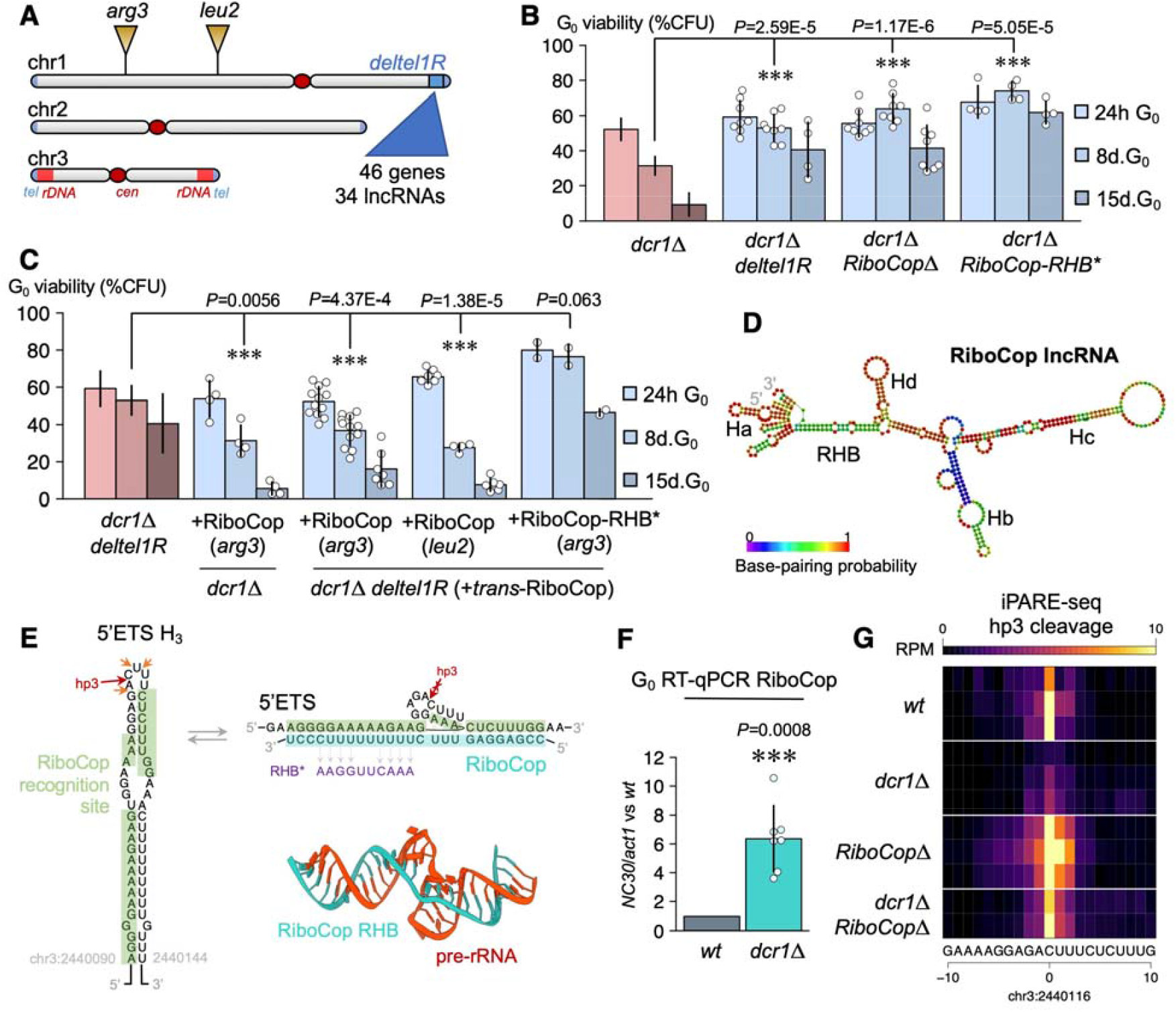
Dicer mutants are rescued by a new trans-acting lncRNA, NC30/RiboCop, involved in pre-rRNA processing regulation. (**A**) Microevolution of the *dcr1*Δ strain resulted in recovery of the suppressor *dcr1*Δ *deltel1R*, harboring a deletion (∼160kb near the right subtelomeric region of chr1). (**B**) The loss of viability during G_0_ maintenance in *dcr1*Δ mutants is suppressed to the same extent in *dcr1*Δ *deltel1R* and *dcr1*Δ*RiboCop*Δ, showing that the lncRNA RiboCop within *deltel1R* is responsible for phenotypic suppression. RiboCop harbors a sequence (RHB) which can pair to the pre-rRNA; RiboCop-RHB* mutants also rescue *dcr1*Δ. (**C**) Re-insertion of RiboCop (including native promoter and terminator) at other intergenic intervals in chr1 in the *dcr1*Δ *deltel1R* suppressor results in the re-acquisition of a *dcr1*Δ phenotype, i.e. anti-suppression. Re-insertion of RiboCop-RHB* does not display this effect, showing that RiboCop within deltel1R is necessary and sufficient for the *dcr1*Δ phenotype and requires its RHB pre-rRNA binding sequence. (**D**) Predicted RiboCop RNA secondary structure (RNAfold and Mfold), showing its 4 main hairpins (Ha-Hd) and RHB region. (**E**) The RiboCop RHB pairs with the 5’ETS third hairpin, containing the new hp3 cleavage site, which is predicted to only be exposed when RiboCop is unpaired. Predicted tertiary structure of RiboCop RHB recognition to the pre-rRNA (AlphaFold3). The sequence of the mutated RHB* is provided. (**F**) While wild-type G0 cells do not express RiboCop, *dcr1*Δ mutants result in >6-fold activation. (**G**) The hp3 cleavage site is inhibited in *dcr1*Δ mutants, and up-regulated in *RiboCop*Δ and *dcr1*Δ*RiboCop*Δ mutants, compatible with a role for hp3 cleavage as a pre-rRNA processing quality control step. (*** represents *P*<0.01, t-test).

Dicer-dependent lncRNAs were strongly enriched for centromeric non-coding RNAs as expected (fig. S3EF), and Dicer-dependent small RNAs were detected from these regions in G_0_ (fig. S3F), although to a lower extent than in S-phase *(15,42)*. Other Dicer-regulated transcripts with G_0_ small RNA were the telomeric helicase gene *tlh1* and the adjacent SPAC212.06c gene, which have homology to *dg/dh* repeats *(43)*. In contrast, we did not detect Dicer-dependent small RNAs originating from the other non-coding RNA loci upregulated in *dcr1*Δ, including NC30 (fig. S3G). Consistent with this absence of siRNAs, none of these lncRNAs displayed an iPARE-seq signal indicative of being Dicer targets, suggesting that they are indirect targets, potentially resulting from the nucleolar stress caused by pre-rRNA processing defects.

### RiboCop is a trans-acting non-coding RNA with snoRNA-like features

Analysis of the regulatory regions of the NC30 locus revealed that it harbors a promoter comprising both a canonical TATA box at position -50 and a HomolD box at position -20 (fig. S3H). The HomolD box (CAGTCACA) is a motif commonly found in the promoter of TATA-less ribosomal protein genes, several snoRNAs, and other housekeeping genes *(44)*, and binds Rrn7, a core factor of RNA polymerase I, which in this context forms a pre-initiation complex for RNA polymerase II *(44,45)*. The dual presence of a HomolD box and a TATA-box is found at the promoter of several lncRNAs, including the U3 snoRNA, where this arrangement was proposed to reflect its special role in ribosomal processing *(46)*, as well as in *nc-tgp1 (47)* and *prt2 (48)*. Given these features and its genetic suppression of Dicer rRNA defects (see below), we hypothesized that NC30 might be involved in rRNA processing and named it “RiboCop” for Ribosomal RNA Co-processor.

According to our hypothesis, we looked for potential regions of complementarity between Dicer-dependent lncRNAs and rRNA (fig. S3I). We found that RiboCop uniquely harbors a 14nt sequence complementary to the start of the 5’ETS pre-rRNA, and modelling this interaction by both RNA:RNA interaction prediction algorithms (DuplexFold, INTARNA) and AI-enabled 3D prediction (AlphaFold3) revealed a 25nt-long recognition sequence, longer than several bona fide U3 snoRNA:pre-rRNA interaction sites *(49,50)*. We termed this sequence in RiboCop the ‘rDNA homology box’ (RHB) (Fig. 2D). To assay whether this region is important for RiboCop function, we created a RiboCop-RHB* mutant (8 SNPs on the 14nt seed RHB sequence) and assayed its ability to rescue a *dcr1*Δ mutant in G_0_. The *dcr1*Δ *RiboCop-RHB\** double-mutant displayed suppression similar to that of *dcr1*Δ*RiboCop*Δ and *dcr1*Δ*deltel1R* strains (Fig. 2BC), indicating that the RHB motif is essential for RiboCop function and strongly suggesting that this function is mediated by pre-rRNA binding. Likewise, *RiboCop-RHB*\* also suppressed *rdp1*Δ and *ago1*Δ (fig. S3K). Overall, these results suggest that RiboCop may be a snoRNA-like ncRNA regulated by Dicer and essential for its phenotypic defects in G_0_ cells.

Suppression of the *dcr1*Δ defect in *dcr1*Δ*deltel1R* and *dcr1*Δ*NC30*Δ mutants suggests that the expression of RiboCop is toxic in *dcr1*Δ cells, and sequence complementarity to pre-rRNA suggests that this effect is exerted in *trans* rather than in *cis*. To test this possibility, we reasoned that re-introducing RiboCop at a different genomic location in the *dcr1*Δ*deltel1R* strain should cancel the *deltel1R* suppression. We re-introduced RiboCop with its endogenous promoter and terminator in the *deltel1R* strain at two distinct genomic locations, near *arg3* and *leu2* respectively (see Methods for exact position). Indeed, these strains displayed a strong quiescence viability defect, similar to the *dcr1*Δ phenotype, showing that RiboCop is both necessary and sufficient for the *dcr1*Δ phenotype (Fig. 2C). At the *arg3* locus, we repeated the *trans*-experiment using the *RiboCop-RHB\** sequence, where the RHB motif in RiboCop is mutated (see above), and the resulting *arg3::NC30-RHB* dcr1*Δ*deltel1R* strain did not result in the *dcr1*Δ phenotype (Fig. 2D). This further confirms that the RHB motif is important for the RiboCop toxic *trans*-effect in the absence of Dicer.

To directly test for a role in pre-rRNA processing, and we performed iPARE-seq in *RiboCop*Δ and *dcr1*Δ*RiboCop*Δ quiescent cells. The only differential iPARE-seq peak in *dcr1*Δ*NC30*Δ corresponded exactly to the predicted binding site at the RHB, reflecting a cleavage site at the tip of the third 5’ETS hairpin in WT cells which we therefore named hp3 (Fig. 2E). Thus, in the absence of Dicer, RiboCop was strongly upregulated (Fig. 2F) and suppressed hp3 cleavage (Fig. 2G). Other Dicer-dependent cleavage sites were not restored, indicating that the suppression by loss of RiboCop occurs after pre-rRNA processing but before rDNA silencing; therefore, the most likely hypothesis is that RiboCop is expressed (or stabilized) as a consequence of the *dcr1*Δ pre-rRNA processing defect, and pauses the earliest steps of pre-rRNA transcription and processing. Attempts to drive expression of RiboCop in G_0_ using various promoters (*p*.*urg1, p*.*urg3, p*.*rpl23, p*.*snu3*) did not result in overexpression (data not shown), indicating that RiboCop RNA was likely stabilized by association with unprocessed rRNA (Fig. 2E). Consistently, RiboCop only accumulates in Dicer mutants (Fig 2F) and other mutants affecting pre-rRNA processing (*rrp6*Δ, see below). In accordance with a tight regulation in quiescence, the RiboCop promoter strongly binds the Clr6 I’ repressive complex in cycling cells *(51)*, and RiboCop is only expressed during late meiosis/sporulation *(11,52)* and in dormant spores *(53)* (fig. S5). Taken together, these results indicate that RiboCop is the key mediator of the quiescence defects of RNAi mutants, and is able to act in *trans*.

### RiboCop mediates G_0_-induced rDNA silencing via the RENT complex

Interestingly, the position of hp3 mirrors that of site A’/01 in mouse and human pre-rRNA (third hairpin from +1 transcription start site) *(54)*, whose inhibition was recently shown to pause RNA pol I transcription during nucleolar stress *(55)*. Consistent with a role in rDNA silencing, we have previously shown that the cause of viability defects in *dcr1*Δ mutants is over-accumulation of H3K9me heterochromatin at the rDNA *(15,18,56)*. To assess at which stage the *dcr1*Δ defects were suppressed by *RiboCop*Δ, we performed ChIP-qPCR of H3K9me2. In *dcr1*Δ cells, H3K9me2 levels are strongly increased in G_0_ (Fig. 3A) and are responsible for the loss of viability *(15)*; indeed, *dcr1*Δ*RiboCop*Δ double-mutants fully suppress the increase in H3K9me2, which returns to the same level as in *RiboCop*Δ or in wild-type cells (Fig 3A). In contrast, *RiboCop*Δ did not affect H3K9me2 levels at pericentromeric heterochromatin (*dg/dh* repeats, using the *otr1R::ade6+ imr1L::ura4+* reporter (Fig. 3B), nor did it affect centromeric silencing assessed by TBZ hypersensitivity, in contrast to *clr4*Δ) (Fig. 3C), but showed a small but statistically significant decrease of H3K9me2 at rDNA in wild-type G_0_ cells (Fig. 3A). These results indicate that RiboCop mediates the G_0_-induced increase of H3K9me2 at rDNA chromatin.

**Fig. 3.**
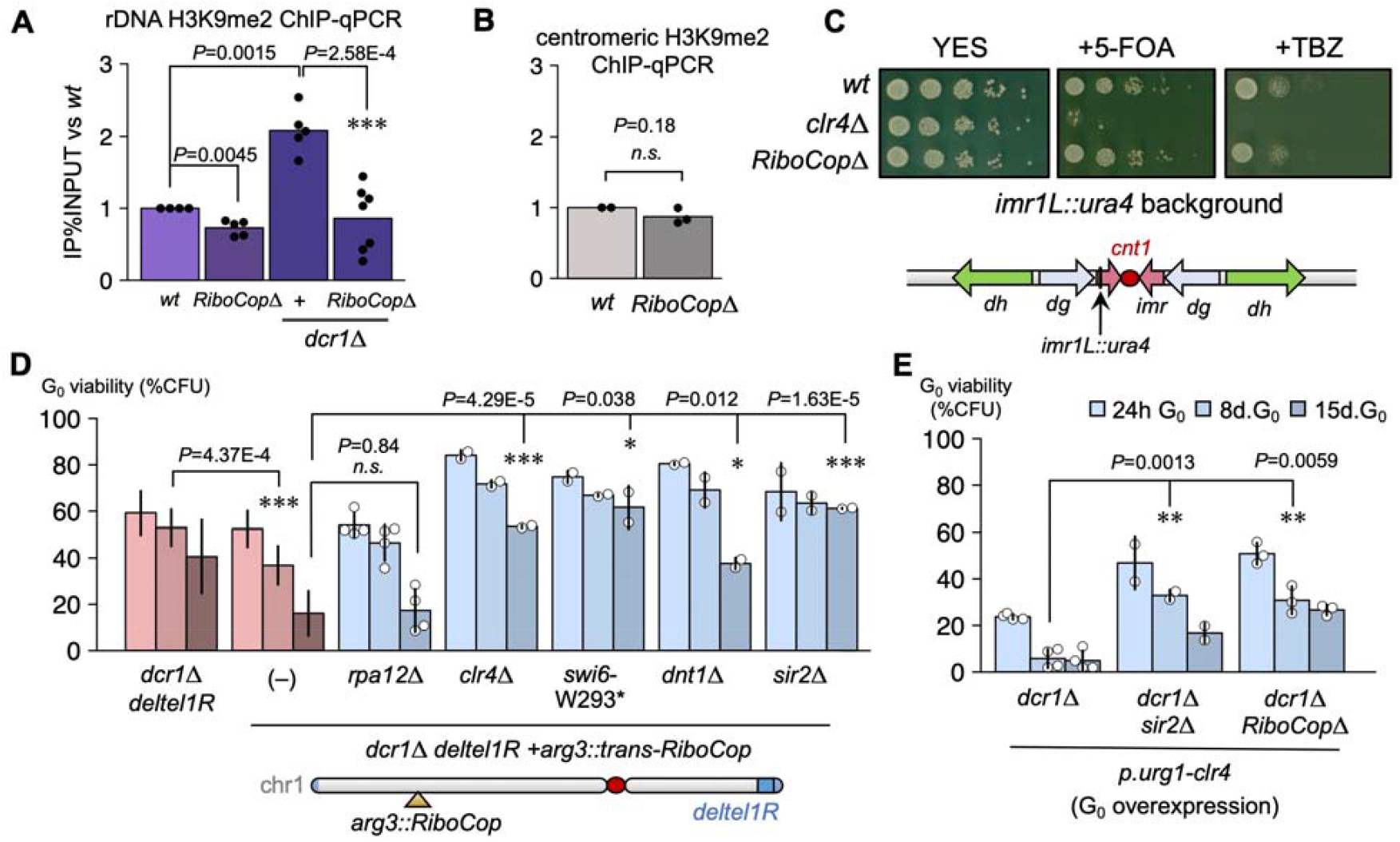
RiboCop triggers rDNA silencing in quiescence. (**A**) H3K9me2 ChIP-qPCR on the rDNA (18S) shows the accumulation of rDNA H3K9me2 in *dcr1*Δ mutants in G_0_. This accumulation is fully suppressed in the *dcr1*Δ*RiboCop*Δ double-mutant. (**B**) *RiboCop*Δ does not affect centromeric H3K9me2 in G_0_ cells, nor (**C**) centromeric silencing in cycling cells, in contrast to the H3K9-methyltransferase mutant *clr4*Δ, as shown by using the *otr1R::ura4+* background (*ura4* de-repression results in 5-FOA sensitivity) and by TBZ resistance (loss of centromeric H3K9me results in TBZ hypersensitivity). (**D**) Analysis of other *dcr1*Δ suppressors in relationship to RiboCop, using the *dcr1*Δ deltel1R arg3::trans-RiboCop phenotypic strain. Suppressors upstream of RiboCop activation display the *dcr1*Δ phenotype of G_0_ viability loss, such as *rpa12*Δ, whereas suppressors downstream of RiboCop activation result in phenotypic suppression, such as *clr4*Δ (H3K9 methyltransferase), *swi6*-W293* (HP1), *dnt1*Δ (RENT rDNA silencing complex) and *sir2*Δ (H3K9 deacetylase). These results are compatible with the hypothesis that RiboCop activation results in rDNA-specific H3K9me silencing. (**E**) Both *RiboCop*Δ and *sir2*Δ result in partial suppression of the *dcr1*Δ *p*.*urg1-clr4* strain, in which rDNA silencing is over-activated^14^. (*: *P*<0.05; **: *P*<0.01; ***: *P*<0.001, t-test).

This raised the question of how a non-coding RNA recruits Clr4 to rDNA? In *S. cerevisiae*, the RENT complex is a nucleolar silencing complex comprised of NET1, SIR2 and CDC14, and these proteins are conserved in *S. pombe*. Moreover, Sir2 has H3K9 deacetylase activity, which mediates the first step in removing H3K9ac to allow methylation to H3K9me2 by Clr4 *(57)*. In *S. cerevisiae*, SIR2 is a key factor in triggering rDNA silencing following nitrogen starvation or rapamycin treatment, conditions which trigger G_0_ *(58)*, and in mammals this function is harbored by sirtuin homologs such as SIRT1 *(59)* and SIRT7 *(60)*. Intriguingly, NET1, while a silencing factor, binds to active rDNA chromatin and to RNA pol I *(61)* and has an activating domain *(62)*, therefore displaying a bivalent function; furthermore, NET1 and SIR2 physically interact *(63,64)*. This raises the possibility that RENT functions as a silencing trigger *(61,63)*, similarly to our proposed role for RiboCop upon nucleolar stress. We therefore hypothesized that the *S. pombe* NET1 ortholog Dnt1 plays a similar role to initiate silencing by

the H3K9 methylation pathway by recruiting Sir2. Consistently, we found that *dcr1*Δ*dnt1*Δ double-mutants displayed complete suppression of the quiescence maintenance defect of *dcr1*Δ (fig. S4) as did a *dcr1*Δ*sir2*Δ double-mutant (fig. S4), which were phenotypically identical to the suppression seen in *dcr1*Δ*clr4*Δ *(15)* and also suppressed *rdp1*Δ and *ago1*Δ (fig. S4). These results suggest that the silencing seen in quiescence in the *dcr1*Δ strain is indeed dependent on the *S. pombe* equivalent to the RENT complex: Dnt1 and Sir2.

We took advantage of expression of NC30/RiboCop in *trans* to place other *dcr1*Δ G_0_ suppressors upstream and downstream of its activity. Suppressors downstream of RiboCop should restore viability to a *arg3::NC30 dcr1*Δ *deltel1R* strain, while suppressors upstream should not, allowing us to identify the genetic requirements for RiboCop-mediated silencing. We found that *dnt1*Δ, *sir2*Δ, *clr4*Δ and *swi6-W293\** all continued to suppress the *dcr1*Δ phenotype when RiboCop was overexpressed, while *rpa12*Δ did not (Fig. 3D), indicating that RiboCop acts downstream of RNA pol I stalling, but upstream of silencing mediated by the RENT complex.

Taken together, these results suggest that RiboCop silences rDNA by recruiting the RENT complex, comprising Dnt1 and the H3K9 deacetylase Sir2, allowing the recruitment of Clr4 and the H3K9 methylation pathway. We previously showed that overexpression of Clr4 in G_0_ cells using *p*.*urg1-clr4* worsens the viability defects of *dcr1*Δ cells by further increasing rDNA heterochromatin formation *(15)*. We found that deleting RiboCop or Sir2 in the *p*.*urg1-clr4 dcr1*Δ background results in phenotypic suppression (Fig. 3E), confirming that RiboCop co-operates with RENT to silence rDNA.

Can nucleolar stress trigger RiboCop-mediated rDNA silencing by RENT independently of a *dcr1*Δ mutant background? In principle, the persistence of uncleaved 5’ETS could be sufficient to trigger this response. We tested this model independently of RNAi via the RNA exosome, whose primary function is pre-rRNA processing *(65)*. The key exosome component Rrp6 (Required for Ribosomal Processing 6) is required for 5.8S processing in *S. cerevisiae (66)*, and for the degradation of the 5’ETS pre-rRNA transcript in yeast, mice and humans *(67,68)*. In budding yeast, the exosome is targeted to pre-rRNA via specific nucleolar proteins such as Utp18 and Nop53^PICT1^ *(68)* and Rrp6 also binds to the 5’ETS pre-rRNA directly *(69)*. In human cells, RRP6 degrades the 5’ETS fragment after A’/01 cleavage; we reasoned that an *rrp6*Δ mutant, like *dcr1*Δ, would similarly result in processing defects downstream of hp3. We therefore assayed the phenotype of the *rrp6*Δ mutant in G_0_, and found that it has strong defects in quiescence maintenance (fig. S5). These maintenance defects were similar to those of RNAi mutants, except that *rrp6*Δ did not have a G_0_-entry defect, which is due to loss of centromeric heterochromatin in RNAi mutants *(15)*. Consistent with our model, RiboCop was strongly induced in *rrp6*Δ G_0_ cells (fig. S5), and *RiboCop*Δ partially restored viability to *rrp6*Δ cells in G_0_ (27% viability vs 14.7% at 8d G_0_ ; p-value<0.02, t-test) (fig. S5). Given that the exosome regulates thousands of ncRNAs *(11)*, the partial suppression of *rrp6*Δ by deleting a single lncRNA is notable. Similar suppression was observed in *rrp6*Δ*sir2*Δ (fig. S5). Overall, these results are in accordance with our model, where RiboCop is activated when pre-rRNA processing fails, likely through stabilization of binding to the 5’ETS via its RHB motif, and recruits the RENT complex and H3K9 methyltransferase Clr4, silencing rDNA and resulting in cell death in G_0_ in RNAi and exosome mutants.

### Identification of the RiboCop riboprotein complex

To understand how RiboCop recruits the RENT complex, we attempted to purify RiboCop-associated proteins. The very low expression level of RiboCop did not allow recovery of the native endogenous complex, even in *dcr1*Δ mutants; instead, we used *in vitro*-transcribed RiboCop (Fig. 4AB), which was 3’-biotinylated and folded. RiboCop was incubated with lysate from *dcr1*Δ cells, and used for pull-down using magnetic streptavidin beads to purify associated proteins (Fig. 4A & Material & Methods). This approach enriches for potential RiboCop interactors depending on their concentration in cellular extracts (i.e. whether they are present in a limiting fashion or in excess). We were able to obtain a clear enrichment of specific proteins in pull-downs in the presence of RiboCop, compared to pull-downs without addition of ncRNA, as imaged by Coomassie Blue (Fig. 4C). Mass-spectrometry analysis of excised bands revealed the presence of RNase H1, Enp2, Fib1 and Nop56, along with several other rRNA processing-related proteins (Kri1, Nop58, Nop4) (Fig 4). We validated these proteins by repeating the purification process and Western blots following pull-down in tagged strain backgrounds (*rnh1*-(Gly)_6_-3xFLAG and *enp2*-(Gly)_6_-3xFLAG) (Fig. 4D). Moreover, we repeated the pulldown in *dcr1*Δ*RiboCop*Δ *rnh1*-(Gly)_6_-3xFLAG cells, and found that RiboCop was still able to interact with RNase H1, indicating this association was not due to artefactual formation to RNA-DNA hybrids with genomic DNA at the native RiboCop locus (Fig. 4D). In addition to RNase H1 and Enp2, several specific nucleolar proteins were identified, including Nop56, Nop58 and fibrillarin (Fib1), which are typically associated with snoRNAs. RiboCop does not have C/D-boxes and snoRNA structure, but it does have a D-box adjacent to the RHB domain; it is therefore not clear whether it constitutes a non-canonical snoRNA-like ncRNA, or if these proteins are recovered indirectly from the U3 snoRNP complex via Enp2. However, the absence of the main U3 snoRNP proteins in the pull-down (UTP-A, UTP-B, UTP-C) suggests specific recruitment of Nop56/Nop58/Fib1; furthermore, Fib1 mutants in human cells have defective processing of A’/01, the proposed equivalent of RiboCop target hp3 *(70)*.

**Fig. 4.**
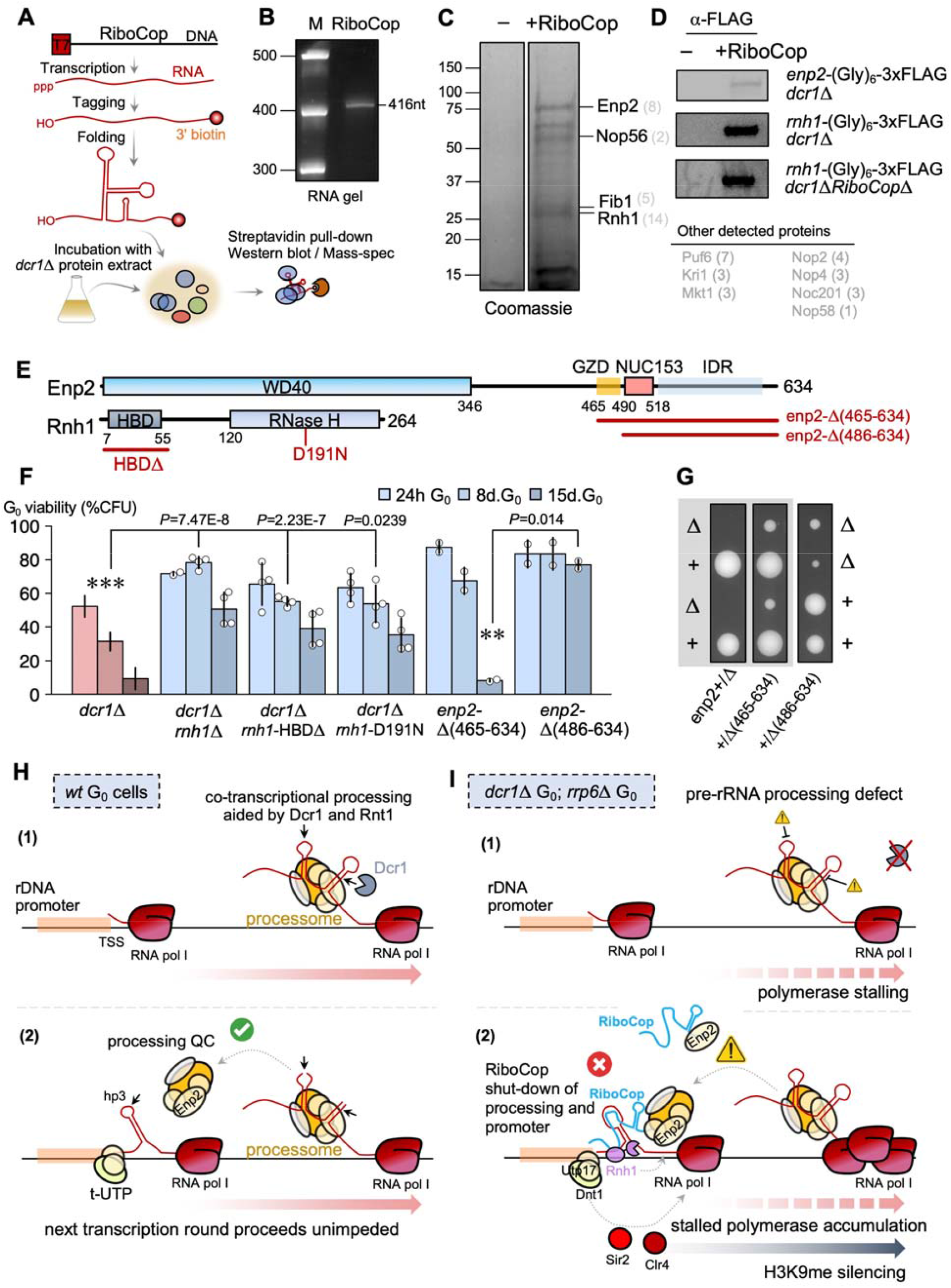
RiboCop forms a riboprotein complex for pre-rRNA processing quality control. (**A**) Workflow for purification of RiboCop-associated proteins: *in vitro*-transcribed, biotinylated and folded RiboCop RNA is purified and incubated with a cell lysate extract from *dcr1*Δ cells. The formed complexes are purified on streptavidin beads and characterized by SDS-PAGE followed by mass-spectrometry (LC/MS) or Western blotting. (**B**) *In vitro*-transcribed RiboCop purity as verified by migration on a 10% TBE-urea RNA gel. (**C**) RiboCop associates with a set of ribosomal processing proteins and RNase H1 (Rnh1). Individual bands from the Coomassie-stained SDS-PAGE gel were excised and identified by MS. (**D**) The entire purification was repeated in tagged strains to confirm that RiboCop pulls down Enp2 and Rnh1. The Rnh1 experiment was repeated in a *dcr1*Δ*RiboCop*Δ background to ensure no artefactual RNA:DNA hybrid formation by excess RiboCop binding at its genetic locus. (**E**) Domain structure organization of Enp2 and Rnh1 proteins. The GZD (G-Zero Defective) is a low-complexity region we identified to be required for G_0_ viability. (*: *P*<0.05, t-test). (**F**) *rnh1*Δ is a suppressor of the *dcr1*Δ G_0_ phenotype; both the HBD and RNase H domains are required, as shown by the suppression in *dcr1*Δ*rnh1-*HBDΔ and *dcr1*Δ*rnh1-*D191N (catalytic-dead) mutants. The GZD domain of Enp2 is required for G_0_ viability. (**G**) Tetrad analyses show that *enp2* is an essential gene (no *enp2*Δ progeny recovered from sporulating a heterozygous *enp2*Δ/+ ade6-210/216 diploid), and its C-terminal deletions are viable. The slow-growth phenotype is caused by the loss of the NUC153 domain. (**H**) Proposed model for wild-type G_0_ cells: in quiescence, RNA pol I transcription requires co-transcriptional processing by the processome, including the RNase III enzymes Dcr1 and Rnt1, in particular at the A_0_ site. Proper processing quality check allows the next round of RNA pol I transcription to proceed, by back-signaling to promoter-associated processome proteins (t-UTP complex: transcriptional U3 RNP-associated) and allowing hp3 cleavage. (**I**) In mutants defective for pre-rRNA processing in quiescence—such as *dcr1*Δ and *rrp6*Δ—the quality check fails. Enp2 recruits and stabilizes RiboCop to the rDNA promoter, where RiboCop inhibits hp3 processing by base-pairing and occluding the cleavage site, and recruits RNase H to degrade the RNA:DNA hybrid formed by RNA pol I transcription and stalling. The blocked rDNA promoter results in the RENT rDNA silencing complex recruiting Sir2 (H3K9 deacetylase) and Clr4 (H3K9 methyltransferase) for silencing the rDNA repeat, following R-loop removal, by H3K9me heterochromatin.

Enp2 is a conserved nucleolar protein in the U3 snoRNP complex *(71)* which guides A_0_ processing *(28)*, and which also interacts with Dnt1 via Utp17(=Nan1) *(64)*, providing a likely pathway for the recruitment of the RENT complex by RiboCop at the rDNA promoter. Enp2 is recruited late to the processome and may play a quality-control role for 18S pre-processing *(71)*. In accordance with this, Enp2^NOL10^ is induced and required for the response to nucleolar stress in human cells *(72)*. In *S. pombe*, Enp2 is also enriched in Swi6^HP1^-associated chromatin *(73)*, making it a likely candidate to bridge processing and silencing machineries in general. Because Enp2 is essential in *S. pombe*, we could not assess whether a deletion mutant modifies the *dcr1*Δ G_0_ phenotype. The C-terminal domain of Enp2 is disordered and is not present in U3 snoRNP structures *(74)*. We generated viable Enp2 C-terminal truncations; *enp2*-E521Δ and *enp2*-P486Δ had wild-type viability while the *enp2*-L465Δ deletion displayed a loss of viability in quiescence, suggesting that the short IDR between the WD40 domain and the NUC153 domain is important for maintaining proper G_0_ function. We termed this region the GZD domain (G Zero Defective). While the combination of the viable alleles *enp2*-E521Δ and *enp2*-P486Δ with *dcr1*Δ did not modify the *dcr1*Δ phenotype, we were unable to recover a viable *enp2*-L465Δ*dcr1*Δ double-mutant, suggesting negative interaction even in cycling cells.

### RNase H1 is required for G0 defects in the absence of Dicer

R-loops accumulate strongly at rDNA in cycling cells of *dcr1*Δ *(75)*, and in mammalian cells RNase H1 is dynamically induced in response to RNA pol I R-loops *(76)*. Therefore, the interaction of RiboCop with RNase H1 could suggest a scenario where the first step towards recruiting rDNA silencing factors is to remove the R-loop left by stalled RNA polymerase I in Dicer mutants. If this model is correct, then loss-of-function of RNase H1 mutants should phenocopy *NC30*Δ in suppressing *dcr1*Δ G_0_ defects. We therefore constructed the single-mutant *rnh1*Δ strain and found that it was viable in G_0_ and was indeed a strong suppressor of *dcr1*Δ (Fig. 4F). Next, we constructed the catalytic mutant Rnh1-D191N analogous to the well-characterized *Bacillus halodurans* D132N mutation *(77)*. Eukaryotic RNase H1 enzymes also contain a N-terminal domain that can bind both RNA:RNA and RNA:DNA hybrids, called the HBD domain (Hybrid Binding Domain) (Fig. 4E). Therefore, we also constructed a mutant strain where this domain was deleted, Rnh1-HBDΔ (removing aminoacids 8-47). Neither of these mutants had a G_0_ defect, and both resulted in strong suppression of the viability loss of *dcr1*Δ in quiescence (Fig. 4F). This suggests that RNase H1 needs to bind a hybrid substrate using its HBD domain, and to cleave an R-loop using its catalytic domain, in the rDNA silencing pathway responsible for *dcr1*Δ G_0_ defects. Overall, these results confirm a genetic interaction between Dicer, RiboCop and RNase H1, and strongly suggest that a key primary step triggering rDNA silencing by Clr4 and Sir2 is to recruit RNase H1 to remove the R-loop resulting from stalled RNA polymerase I.

### Model for nucleolar RNAi and conservation of the nucleolar role of Dicer

In conclusion, we demonstrate that the RNase III family Dicer and Rnt1 nucleases have conserved roles in ribosomal RNA pre-processing, by cleaving multiple sites in the 5’ETS (including the key A_0_ site) and 3’ETS during pre-rRNA transcription, particularly in G_0_ when rRNA processing is predominantly co-transcriptional. In the absence of Dicer, rRNA processing defects result in the activation of RiboCop by stabilizing the lncRNA, via Enp2 quality control of processome activity. RiboCop binds to the unprocessed 5’ETS proximal to the rDNA promoter, to pause RNA pol I transcription, analogously to nucleolar stress in human cells *(52)*. At the rDNA promoter, RiboCop triggers silencing of the rDNA repeat by (i) recruiting RNase H to degrade R-loops following the last round of RNA pol I transcription, and (ii) recruiting the RENT complex (Dnt1, Sir2) and Clr4 via the U3 snoRNP (Utp17), resulting in H3K9 methylation and rDNA heterochromatin formation. Transcriptomic analyses show that RiboCop is only expressed in wild-type cells at the end of meiosis when sporulation starts *(11,54)* and in dormant spores *(55)* (fig. S5), suggesting that its physiological role is related to cellular dormancy (where rDNA would become completely silent). This is compatible with our proposed function for RiboCop in inhibiting the earliest processing sites of the pre-rRNA, and triggering rDNA silencing via recruitment of RNase H, processing inhibition, and recruitment of H3K9 methylation. Processing mutants, such as *dcr1*Δ and *rrp6*Δ in quiescent cells (maintaining metabolic activity), may therefore abnormally trigger the complete rRNA shut-down as in dormant cells (non-metabolic spores) via the spurious activation of the RiboCop/Enp2/Rnh1 RNP.

Using ChIP-exo, we have been able to show direct Dicer binding to rDNA chromatin consistent with observations in cycling cells using DamID *(40)*, and in mammalian cells *(41)*. In fact, other RNAi proteins are also nucleolar, like AGO2 which is recruited to rRNA in a Dicer-dependent manner *(78)* and the Microprocessor complex which physically associates with nucleolin *(79)*, or the recovery of RNA-dependent RNA polymerase activity in the cauliflower U3 snoRNP *(80)*. Moreover, small RNAs derived from rRNA—an indirect signature of Dicer— have been detected in a number of species, from cycling *S. pombe* cells at the 5’ETS and 3’ETS (including *priRNA1*) *(15,37,81)* to *Neurospora* (qiRNA), zebrafish, *Drosophila* and mouse *(82)*. Other pre-rRNA associated RNAs, such as the U3 snoRNA, have also been found to be targeted by Dicer in human cells, between the A/A’ and C/D boxes (i.e. the region that forms dsRNA with pre-rRNA) *(82)*, and a 5.8S processing defect was seen upon knockdown of Dicer or Ago2 in HeLa cells *(83)*, although this study did not examine processing of the 5’ETS. Another parallel was found in a recent study showing that RNAi regulates ribosomal protein genes in *Aspergillus fumigatus* conidia (quiescent asexual spores) independently of small RNA *(84)*. Dicer is a member of the RNase III family of nucleases, and Rnt1 is also involved in ribosomal RNA processing in *S. pombe (85)*, and *S. cerevisiae (86)*. In fact, this conservation may be very ancient as one of the main targets of bacterial RNase III is the ribosomal RNA operon *(87)*. In the pathogenic yeast *Cryptococcus neoformans*, a recent study identified several factors involved in RNAi and transposon suppression, which included another RNase III enzyme as well as nucleolar proteins copurifying with fibrillarin and several U3 snoRNP-associated proteins *(88)*. Therefore, our finding of an essential nucleolar function of Dicer in *S. pombe* and of Dicer-dependent rRNA cleavage sites suggests that ribosomal processing is an ancestral function of all RNase III enzymes. In this regard, it is interesting that the A_0_ site is thought to be RNA-directed (by the U3 snoRNA box A/A’), drawing a parallel to other types of RNA-directed RNA cleavage directed by RNAi.

### Is the NC30/RiboCop complex conserved?

We did not identify NC30 orthologs at the sequence level in other fission yeasts (*Schizosaccharomyces* spp.) nor in other members of the subphylum Taphrinomycotina. However, it is important to note that the external and internal spacer elements (ETS/ITS) are also not conserved at the sequence level, and therefore any pre-rRNA processing-associated lncRNA would co-evolve with its sequence. Despite the lack of sequence conservation, the 5’ETS pre-rRNA shows a structural organization consisting of a succession of long hairpins in most species, ranging from budding yeast *(89,90), S. pombe (35,89)*, mouse *(29,91)* and humans *(92)*. We showed that RiboCop protects against cleavage at its NC30:5’ETS binding site on the third helix (herein, ‘hp3 cleavage site’), which has an equivalent site on the third helix of the 5’ETS, the first pre-rRNA cleavage site (the A’/01 site) whose inhibition is essential to maintain nucleolar integrity in mammals *(52)*. Enp2/NOL10 is likewise required for nucleolar integrity during the nucleolar stress response *(72)*, and Fib1 is required for A’/01 processing *(70)*. It is therefore tempting to speculate that a functional equivalent to NC30/RiboCop may exist in mammals, with a diverged RNA sequence, assisting in A’/01 cleavage inhibition during nucleolar stress and/or dormancy.

In mammals, rDNA silencing relies on non-coding promoter RNA (pRNA) in association with the NoRC complex *(20)*. While a key difference between NC30 and pRNA is that pRNA is transcribed in *cis* while NC30 acts in *trans*, these different complexes may converge in their rDNA silencing function, as NoRC recruits the Sir2 ortholog SIRT7 in mammalian cells *(60)*, and RiboCop also functions via Sir2 (SupFig 8B). However pRNA itself, and the pRNA-associated protein TIP5, are specific to mammals. A recent study on a patient-derived pleuropulmonary blastoma model with DICER1 hotspot mutations found that it was greatly sensitive to RNA pol I inhibitors such as CX-5461 (pidnarulex) *(93)*, in accordance with our proposed model of Dicer’s tumorigenicity being caused by its nucleolar function rather than miRNA dysregulation *(94)*. Furthermore, Dicer is known to target several pre-rRNA-associated snoRNAs in mammals, such as the U3 snoRNA involved in A_0_ cleavage *(83,95)*. Enp2/NOL10 expression had a significant effect in prostate cancer prognosis and severity *(96)*; NOL10 is frequently mutated in specific cancers (4% in endometrial cancers cf. BioPortal) and is a critical dependency in some acute myeloid leukemias *(97)*. Further studies on ribosomal quality control by non-coding RNAs during nucleolar stress will therefore likely open many new avenues that could yield therapeutic approaches not only in DICER1 syndrome patients, but potentially also other cancers linked to genetic mutations in RNA surveillance machineries.

## Acknowledgments

The authors would like to thank the fission yeast database PomBase (http://www.pombase.org), which has been essential in the course of this work, including and not limited to: obtaining up-to-date annotations for the pombe genome, exploring genetic interactions and cross-referenced data between sets of genes. The authors would also like to thank the *S. pombe* community for insightful discussion.

## Author contributions

Conceptualization: BR, RAM

Methodology, Investigation: BR

Funding acquisition: RAM

Writing – original draft: BR

Writing – review & editing: BR, RAM

## Data and materials availability

All yeast strains generated in this study are provided in Table S1 and are avaible upon request to the lead contact. All next-generation sequencing data is available on the NCBI SRA database at the following accession number: BioProject PRJNA1328910.

## Supplementary Materials

Materials and Methods

Figs. S1 to S5

Tables S1 to S2

References (*98-120*)

## References and Notes

1. A. de Morree, T.A. Rando. Regulation of adult stem cell quiescence and its functions in the maintenance of tissue integrity. Nat Rev Mol Cell Biol 24(5):334–354 (2023). doi:10.1038/s41580-022-00568-6

2. C.T.J. van Velthoven, T.A. Rando. Stem cell quiescence: dynamism, restraint, and cellular idling. Cell Stem Cell 24(2):213–225 (2019). doi:10.1016/j.stem.2019.01.001

3. B. Roche, B. Arcangioli, R.A. Martienssen. Transcriptional reprogramming in cellular quiescence. RNA Biol 14(7):843–853 (2017). doi:10.1080/15476286.2017.1327510

4. L.L. Breeden, T. Tsukiyama. Quiescence in Saccharomyces cerevisiae. Annu Rev Genet 56:253–278 (2022). doi:10.1146/annurev-genet-080320-023632

5. S. Marguerat, A. Schmidt, S. Codlin, W. Chen, R. Aebersold, J. Bähler. Quantitative analysis of fission yeast transcriptomes and proteomes in proliferating and quiescent cells. Cell 151(3):671–683 (2012). doi:10.1016/j.cell.2012.09.019

6. C.T.J. van Velthoven, A. de Morree, I.M. Egner, J.O. Brett, T.A. Rando. Transcriptional profiling of quiescent muscle stem cells in vivo. Cell Rep 21(7):1994–2004 (2017). doi:10.1016/j.celrep.2017.10.037

7. A. Brunet, M.A. Goodell, T.A. Rando. Ageing and rejuvenation of tissue stem cells and their niches. Nat Rev Mol Cell Biol 24(1):45–62 (2023). doi:10.1038/s41580-022-00510-w

8. J. Fröbel, T. Landspersky, G. Percin, C. Schreck, S. Rahmig, A. Ori, D. Nowak, M. Essers, C. Waskow, R.A.J. Oostendorp. The hematopoietic bone marrow niche ecosystem. Front Cell Dev Biol 9:705410 (2021). doi:10.3389/fcell.2021.705410

9. S.S. Su, Y. Tanaka, I. Samejima, K. Tanaka, M. Yanagida. A nitrogen starvation-induced dormant G0 state in fission yeast: the establishment from uncommitted G1 state and its delay for return to proliferation. J Cell Sci 109(6):1347–1357 (1996). doi:10.1242/jcs.109.6.1347

10. M. Yanagida. Cellular quiescence: are controlling genes conserved? Trends Cell Biol 19(12):705–715 (2009). doi:10.1016/j.tcb.2009.09.006

11. S.R. Atkinson, S. Marguerat, D.A. Bitton, M. Rodríguez-López, C. Rallis, J.F. Lemay, C. Cotobal, M. Malecki, P. Smialowski, J. Mata, P. Korber, F. Bachand, J. Bähler. Long noncoding RNA repertoire and targeting by nuclear exosome, cytoplasmic exonuclease, and RNAi in fission yeast. RNA 24(9):1195–1213 (2018). doi:10.1261/rna.065524.118

12. A. Nevers, A. Doyen, C. Malabat, B. Néron, T. Kergrohen, A. Jacquier, G. Badis. Antisense transcriptional interference mediates condition-specific gene repression in budding yeast. Nucleic Acids Res 46(12):6009–6025 (2018). doi:10.1093/nar/gky342

13. K. Soureas, M.A. Papadimitriou, K. Panoutsopoulou, K.M. Pilala, A. Scorilas, M. Avgeris. Cancer quiescence: non-coding RNAs in the spotlight. Trends Mol Med 29(10):843–858 (2023). doi:10.1016/j.molmed.2023.07.003

14. M. Rodriguez-Lopez, S. Anver, C. Cotobal, S. Kamrad, M. Malecki, C. Correia-Melo, M. Hoti, S.J. Townsend, S. Marguerat, S.K. Pong, M.Y. Wu, L. Montemayor, M. Howell, M. Ralser, J. Bähler. Functional profiling of long intergenic non-coding RNAs in fission yeast. Elife 11:e76000 (2022). doi:10.7554/eLife.76000

15. B. Roche, B. Arcangioli, R. Martienssen. RNA interference is essential for cellular quiescence. Science354(6313):aah5651 (2016). doi:10.1126/science.aah5651

16. K. Sajiki, M. Hatanaka, T. Nakamura, K. Takeda, M. Shimanuki, T. Yoshida, Y. Hanyu, T. Hayashi, Y. Nakaseko, M. Yanagida. Genetic control of cellular quiescence in S. pombe. J Cell Sci 122(9):1418–1429 (2009). doi:10.1242/jcs.046466

17. R.I. Joh, J.S. Khanduja, I.A. Calvo, M. Mistry, C.M. Palmieri, A.J. Savol, S.J.H. Sui, R.I. Sadreyev, M.J. Aryee, M. Motamedi. Survival in quiescence requires the euchromatic deployment of Clr4/SUV39H by Argonaute-associated small RNAs. Mol Cell 64(6):1088–1101 (2016). doi:10.1016/j.molcel.2016.11.020

18. X. Deng, Q. Yao, A. Horvath, Z. Jiang, J. Zhao, T. Fischer, T. Sugiyama. The fission yeast ortholog of Coilin, Mug174, forms Cajal body-like nuclear condensates and is essential for cellular quiescence. Nucleic Acids Res 52(15):9174–9192 (2024). doi:10.1093/nar/gkae463

19. Y. Zahedi, S. Zeng, K. Ekwall. An essential role for the Ino80 chromatin remodeling complex in regulation of gene expression during cellular quiescence. Chromosome Res 31(2):14 (2023). doi:10.1007/s10577-023-09723-x

20. S. Wehner, A.K. Dörrich, P. Ciba, A. Wilde, M. Marz. pRNA: NoRC-associated RNA of rRNA operons. RNA Biol 11(1):3–9 (2014). doi:10.4161/rna.27448

21. C. Ellermeier, E.C. Higuchi, N. Phadnis, L. Holm, J.L. Geelhood, G. Thon, G.R. Smith. RNAi and heterochromatin repress centromeric meiotic recombination. Proc Natl Acad Sci USA 107(19):8701–8705 (2010). doi:10.1073/pnas.0914160107

22. M.A. German, M. Pillay, D.H. Jeong, A. Hetawal, S. Luo, P. Janardhanan, V. Kannan, L.A. Rymarquis, K. Nobuta, R. German, E. De Paoli, C. Lu, G. Schroth, B.C. Meyers, P.J. Green. Global identification of microRNA-target RNA pairs by parallel analysis of RNA ends. Nat Biotechnol 26(8):941–946 (2008). doi:10.1038/nbt1417.

23. G. Rotondo, D. Frendeway. Purification and characterization of the Pac1 ribonuclease of Schizosaccharomyces pombe.Nucleic Acids Res 24(12):2377–2386 (1996). doi:10.1093/nar/24.12.2377

24. J. Gan, J.E. Tropea, B.P. Austin, D.L. Court, D.S. Waugh, X. Ji. Structural insight into the mechanism of double-stranded RNA processing by ribonuclease III. Cell 124(2):355–366 (2006). doi:10.1016/j.cell.2005.11.034

25. D.L. Court, J. Gan, Y.H. Liang, G.X. Shaw, J.E. Tropea, N. Costantino, D.S. Waugh, X. Ji. RNase III: Genetics and function; structure and mechanism. Annu Rev Genet 47:405–431 (2013). doi:10.1146/annurev-genet-110711-155618

26. A.K. Henras, C. Clisson-Chastang, M.F. O’Donohue, A. Chakraborty, P.E. Gleizes. An overview of pre-ribosomal RNA processing in eukaryotes. Wiley Interdiscip Rev RNA 6(2):225–242 (2015). doi:10.1002/wrna.1269

27. E. Ivakine, K. Spasov, D. Frendeway, R.N. Nazar, R.N. Functional significance of intermediate cleavages in the 3’ETS of the pre-rRNA from Schizosaccharomyces pombe.Nucleic Acids Res 31(24):7110–7116 (2003). doi:10.1093/nar/gkg932

28. S.J. Lee, S.J. Baserga. Functional separation of pre-rRNA processing steps revealed by truncation of the U3 small nucleolar ribonucleoprotein component, Mpp10. Proc Natl Acad Sci USA 94(25):13536–13541 (1997). doi:10.1073/pnas.94.25.13536

29. T. Kent, Y.R. Lapik, D.G. Pestov. The 5’ external transcribed spacer in mouse ribosomal RNA contains two cleavage sites. RNA 15(1):14–20 (2009). doi:10.1261/rna.1384709

30. S.A. Elela, H. Igel, M.J. Ares. RNase III cleaves eukaryotic preribosomal RNA at a U3 snoRNP-dependent site. Cell 85(1):115–124 (1996). doi:10.1016/s0092-8674(00)81087-9

31. J. Kufel, B. Dichtl, D. Tollervey. Yeast Rnt1p is required for cleavage of the pre-ribosomal RNA in the 3’ETS but not the 5’ETS. RNA 5(7):909–917 (1999). doi:10.1017/s135583829999026x

32. S. Watt, J. Mata, L. López-Maury, S. Marguerat, G. Burns, J. Bähler. urg1: a uracil-regulatable promoter system for fission yeast with short induction and repression times. PLoS One 3(1):e1428 (2008). doi:10.1371/journal.pone.0001428

33. C.E. Scull, D.A. Schneider. Coordinated control of rRNA processing by RNA polymerase I. Trends Genet 35(10):724–733 (2019). doi:10.1016/j.tig.2019.07.002

34. J. Cheng, N. Kellner, O. Berninghausen, E. Hurt, R. Beckmann. 3.2-Å-resolution structure of the 90S preribosome before A1 pre-rRNA cleavage. Nat Struct Mol Biol 24(11):954–964 (2017). doi:10.1038/nsmb.3476

35. R.V. Intine, L. Good, R.N. Nazar. Essential structural features in the Schizosaccharomyces pombe pre-rRNA 5’ external transcribed spacer. J Mol Biol 286(3):695–708 (1999). doi:10.1006/jmbi.1998.2502

36. L. Bergeron, J.P. Perreault, S.A. Elela, S.A. Short RNA duplexes guide sequence-dependent cleavage by human Dicer. RNA 16(12):2464–2473 (2010). doi:10.1261/rna.2346510

37. R. Jain, N. Iglesias, D. Moazed. Distinct functions of Argonaute Slicer in siRNA maturation and heterochromatin formation. Mol Cell 63(2):191–205 (2016). doi:10.1016/j.molcel.2016.05.039

38. T.A. Volpe, C. Kidner, I.M. Hall, G. Teng, S.I.S. Grewal, R.A. Martienssen. Regulation of heterochromatic silencing and histone H3 lysine-9 methylation by RNAi. Science 297(5588):1833–1837 (2002). doi:10.1126/science.1074973

39. S. Emmerth, H. Schober, D. Gaidatzis, T. Roloff, K. Jacobeit, M. Bühler. Nuclear retention of fission yeast Dicer is a prerequisite for RNAi-mediated heterochromatin assembly. Dev Cell 18(1):102–113 (2010). doi:10.1016/j.devcel.2009.11.011

40. K.J. Woolcock, D. Gaidatzis, T. Punga, M. Bühler. Dicer associates with chromatin to repress genome activity in Schizosaccharomyces pombe. Nat Struct Mol Biol 18(1):94–99 (2011). doi:10.1038/nsmb.1935

41. L. Sinkkonen, T. Hugenschmidt, W. Filipowicz, P. Svoboda, P. Dicer is associated with ribosomal DNA chromatin in mammalian cells. PLoS One 5(8):e12175 (2010). doi:10.1371/journal.pone.0012175

42. A. Kloc, M. Zaratiegui, E. Nora, R. Martienssen. RNA interference guides histone modification during the S phase of chromosomal replication. Curr Biol 18(7):490–495 (2008). doi:10.1016/j.cub.2008.03.016

43. K.R. Hansen, P.T. Ibarra, G. Thon. Evolutionary-conserved telomere-linked helicase genes of fission yeast are repressed by silencing factors, RNAi components and the telomere-binding protein Taz1. Nucleic Acids Res 34(1):78–88 (2006). doi:10.1093/nar/gkj415

44. D.A. Rojas, S. Moreira-Ramos, S. Zock-Emmenthal, F. Urbina, J. Contreras-Levicoy, N.F. Käufer, E. Maldonado. Rrn7 protein, an RNA polymerase I transcription factor, is required for RNA polymerase II-dependent transcription directed by core promoters with a HomolD box sequence. J Biol Chem 286(30):26480–26486 (2011). doi:10.1074/jbc.M111.224337

45. M. Montes, S. Moreira-Ramos, D.A. Rojas, F. Urbina, N.F. Käufer, E. Maldonado. RNA polymerase II components and Rrn7 form a preinitiation complex on the HomolD box to promote ribosomal protein gene expression in Schizosaccharomyces pombe.FEBS J 284(4):615–633 (2017). doi:10.1111/febs.14006

46. S. Nabavi, R.N. Nazar. U3 snoRNA promoter reflects the RNA’s function in ribosome biogenesis. Curr Genet 54(4):175–184 (2008). doi:10.1007/s00294-008-0210-1

47. A.M. Sanchez, S. Shuman, B. Schwer, B. Poly(A) site choice and Pol2 CTD Serine-5 status govern lncRNA control of phosphate-responsive tgp1 gene expression in fission yeast. RNA 24(2):237–250 (2018). doi:10.1261/rna.063966.117

48. A. Garg, A.M. Sanchez, S. Shuman, B. Schwer. A long noncoding (lnc)RNA governs expression of the phosphate transporter Pho84 in fission yeast and has cascading effects on the flanking prt lncRNA and pho1 genes. J Biol Chem 293(12):4456–4467 (2018). doi:10.1074/jbc.RA117.001352

49. L. Good, R.V. Intine, R.N. Nazar. The ribosomal-RNA-processing pathway in Schizosaccharomyces pombe. Eur J Biochem 247(1):314–321 (1997). doi:10.1111/j.1432-1033.1997.00314.x

50. D.A. Samarsky, M.J. Fournier. Functional mapping of the U3 small nucleolar RNA from the yeast Saccharomyces cerevisiae. Mol Cell Biol 18(6):3431–3444 (1998). doi:10.1128/MCB.18.6.3431

51. N. Zilio, S. Codlin, A.A. Vashisht, D.A. Bitton, S.R. Head, J.A. Wohlschlegel, J. Bähler, M.N. Boddy. A novel histone deacetylase complex in the control of transcription and genome stability. Mol Cell Biol 34(18):3500–3514 (2014). doi:10.1128/MCB.00519-14

52. Y. Wei, N.N. Lee, L. Pan, J. Dhakshnamoorthy, L.L. Sun, M. Zofall, D. Wheeler, S.I.S. Grewal. TOR targets an RNA processing network to regulate facultative heterochromatin, developmental gene expression and cell proliferation. Nat Cell Biol 23(3):243–256 (2021). doi:10.1038/s41556-021-00631-y

53. H. Tsuyuzaki, M. Hosokawa, K. Arikawa, T. Yoda, N. Okada, H. Takeyama, M. Sato. Time-lapse single-cell transcriptomics reveals modulation of histone H3 for dormancy breaking in fission yeast. Nat Commun 11(1):1265 (2020). doi:10.1038/s41467-020-15060-y

54. S.T. Mullineux, D.L.J. Lafontaine. Mapping the cleavage sites on mammalian pre-rRNAs: where do we stand? Biochimie 94(7):1521–1532 (2012). doi:10.1016/j.biochi.2012.02.001

55. W. Szaflarski, M. Lesniczak-Staszak, M. Sowinski, S. Ojha, A. Aulas, D. Dave, S. Malla, P. Anderson, P. Ivanov, S.M. Lyons. Early rRNA processing is a stress-dependent regulatory event whose inhibition maintains nucleolar integrity. Nucleic Acids Res 50(2):1033–1051 (2022). doi:10.1093/nar/gkab1231.

56. H. Hirai, N. Takemata, M. Tamura, K. Ohta. Facultative heterochromatin formation in rDNA is essential for cell survival during nutritional starvation. Nucleic Acids Res 50(7):3727–3744 (2022). doi:10.1093/nar/gkac175

57. B.J. Alper, G. Job, R.K. Yadav, S. Shanker, B.R. Lowe, J.F. Partridge. Sir2 is required for Clr4 to initiate centromeric heterochromatin assembly in fission yeast. EMBO J 32(17):2321–2335 (2013). doi:10.1038/emboj.2013.143

58. C.W. Ha, W.K. Huh. The implication of Sir2 in replicative aging and senescence in Saccharomyces cerevisiae. Aging (Albany NY) 3(3):319–324 (2011). doi:10.18632/aging.100299

59. G. Kreiner, A. Sönmez, B. Liss, R. Parlato. Integration of the deacetylase SIRT1 in the response to nucleolar stress: metabolic implications for neurodegenerative diseases. Front Mol Neurosci 12:106 (2019). doi:10.3389/fnmol.2019.00106

60. S. Paredes, M. Angulo-Ibanez, L. Tasselli, S.M. Carlson, W. Zheng, T.M. Li, K.F. Chua. The epigenetic regulator SIRT7 guards against mammalian cellular senescence induced by ribosomal DNA instability. J Biol Chem 293(28):11242–11250 (2018). doi:10.1074/jbc.AC118.003325

61. W. Shou, K.M. Sakamoto, J. Keener, K.W. Morimoto, E.E. Traverso, R. Azzam, G.J. Hoppe, R.M. Feldman, J. DeModena, D. Moazed, H. Charbonneau, M. Nomura, R.J. Deshaies. Net1 stimulates RNA polymerase I transcription and regulates nucleolar structure independently of controlling mitotic exit. Mol Cell 8(1):45–55 (2001). doi:10.1016/s1097-2765(01)00291-x

62. K. Hannig, V. Babl, K. Hergert, A. Maier, M. Pilsl, C. Schächner, U. Stöckl, P. Milkereit, H. Tschochner, W. Seufert, J. Griesenbeck. The C-terminal region of Net1 is an activator of RNA polymerase I transcription with conserved features from yeast to human. PLoS Genet 15(2):e1008006 (2019). doi:10.1371/journal.pgen.1008006

63. W. Shou, J.H. Seol, A. Shevchenko, C. Baskerville, D. Moazed, Z.W. Chen, J. Jang, A. Shevchenko, H. Charbonneau, R.J. Deshaies. Exit from mitosis is triggered by Tem1-dependent release of the protein phosphatase Cdc14 from nucleolar RENT complex. Cell 97(2):233–244 (1999). doi:10.1016/s0092-8674(00)80733-3

64. A.F. Straight, W. Shou, G.J. Dowd, C.W. Turck, R.J. Deshaies, A.D. Johnson, D. Moazed. Net1, a Sir2-associated nucleolar protein required for rDNA silencing and nucleolar integrity. Cell 97(2):245–256 (1999). doi:10.1016/s0092-8674(00)80734-5

65. E.K. Okuda, F.A. Gonzales-Zubiate, O. Gadal, C.C. Oliveira. Nucleolar localization of the yeast RNA exosome subunit Rrp44 hints at early pre-rRNA processing as its main function. J Biol Chem 295(32):11195–11213 (2020). doi:10.1074/jbc.RA120.013589

66. M.W. Briggs, K.T. Burkard, J.S. Butler. Rrp6p, the yeast homologue of the human PM-Scl 100kDa autoantigen, is essential for efficient 5.8S rRNA 3’ end formation. J Biol Chem 273(21):13255–13263 (1998). doi:10.1074/jbc.273.21.13255

67. K. Kobyłecki, K. Drążkowska, T.M. Kuliński, A. Dziembowski, R. Tomecki. Elimination of 01/A’-A0 pre-rRNA processing by-product in human cells involves cooperative action of two nuclear exosome-associated nucleases: RRP6 and DIS3. RNA 24(12):1677–1692 (2018). doi:10.1261/rna.066589.118

68. M. Thoms, E. Thomson, J. Baßler, M. Gnädig, S. Griesel, E. Hurt. The exosome is recruited to RNA substrates through specific adaptor proteins. Cell 162(5):1029–1038 (2015). doi:10.1016/j.cell.2015.07.060

69. C. Delan-Forino, C. Schneider, D. Tollervey. Transcriptome-wide analysis of alternative routes for RNA substrates into the exosome complex. PLoS Genet 13(3):e1006699 (2017). doi:10.1371/journal.pgen.1006699

70. L. Tafforeau, C. Zorbas, J.L. Langhendries, S.T. Mullineux, V. Stamatopoulou, R. Mullier, L. Wacheul, D.L.J. Lafontaine. The complexity of human ribosome biogenesis revealed by systematic nucleolar screening of pre-rRNA processing factors. Mol Cell 51(4):539–551 (2013). doi:10.1016/j.molcel.2013.08.011

71. S. Soltanieh, M. Lapensée, F. Dragon. Nucleolar proteins Bfr2 and Enp2 interact with DEAD-box RNA helicase Dpb4 in two different complexes. Nucleic Acids Res 42(5):3194–3206 (2014). doi:10.1093/nar/gkt1293

72. X. Jin, H. Tanaka, M. Jin, K. Fujita, H. Homma, M. Inotsume, H. Yong, K. Umeda, N. Kodera, T. Ando, H. Okazawa. PQBP5/NOL10 maintains and anchors the nucleolus under physiological and osmotic stress conditions. Nat Commun 14(1):9 (2023). doi:10.1038/s41467-022-35602-w

73. N. Iglesias, J.A. Paulo, A. Tatarkis, X. Wang, A.L. Edwards, N.V. Bhanu, B.A. Garcia, W. Haas, S.P. Gygi, D. Moazed. Native chromatin proteomics reveals a role for specific nucleoporins in heterochromatin organization and maintenance. Mol Cell 77(1):51–66 (2020). doi:10.1016/j.molcel.2019.10.018

74. J. Cheng, J. Baßler, P. Fischer, B. Lau, N. Kellner, R. Kunze, S. Griesel, M. Kallas, O. Berninghausen, D. Strauss, R. Beckmann, E. Hurt. Thermophile 90S pre-ribosome structures reveal the reverse order of co-transcriptional 18S rRNA subdomain integration. Mol Cell 75(6):1256–1269 (2019). doi:10.1016/j.molcel.2019.06.032

75. S.E. Castel, J. Ren, S. Bhattacharjee, A.Y. Chang, M. Sánchez, A. Valbuena, F. Antequera, R.A. Martienssen. Dicer promotes transcription termination at sites of replication stress to maintain genome stability. Cell 159(3):572–583 (2014). doi:10.1016/j.cell.2014.09.031

76. W. Shen, H. Sun, C.L. De Hoyos, J.K. Bailey, X.H. Liang, S.T. Crooke. Dynamic nucleoplasmic and nucleolar localization of mammalian RNase H1 in response to RNAP I transcriptional R-loops. Nucleic Acids Res 45(18):10672–10692 (2017). doi:10.1093/nar/gkx710

77. M. Nowotny, S.A. Gaidamakov, R.J. Crouch, W. Yang. Crystal structures of RNase H bound to an RNA/DNA hybrid: substrate specificity and metal-dependent catalysis. Cell 121(7):1005–1016 (2005). doi:10.1016/j.cell.2005.04.024

78. B.L. Atwood, J.L. Woolnough, G.M. Lefevre, M. Saint Just Ribeiro, G. Felsenfeld, K.E. Giles. Human Argonaute 2 is tethered to ribosomal RNA through microRNA interactions. J Biol Chem 291(34):17919–17928 (2016). doi:10.1074/jbc.M116.725051

79. B.F. Pickering, D. Yu, M.W. Van Dyke. Nucleolin protein interacts with microprocessor complex to affect biogenesis of microRNAs 15a and 16. J Biol Chem 286(51):44095–44103 (2011). doi:10.1074/jbc.M111.265439

80. H. Samaha, V. Delorme, F. Pontvianne, R. Cooke, F. Delalande, A. Van Dorsselaer, M. Echeverria, J. Sáez-Vásquez, J. Identification of protein factors and U3 snoRNAs from a Brassica oleracea RNP complex involved in the processing of pre-rRNA. Plant J 61(3):383–398 (2010). doi:10.1111/j.1365-313X.2009.04061.x

81. M. Lambert, A. Benmoussa, P. Provost. Small non-coding RNAs derived from eukaryotic ribosomal RNA. Noncoding RNA 5(1):16 (2019). doi:10.3390/ncrna5010016

82. N. Lemus-Diaz, R.R. Ferreira, K.E. Bohnsack, J. Gruber, M.T. Bohnsack, M.T. The human box C/D snoRNA U3 is a miRNA source and miR-U3 regulates expression of sortin nexin 27. Nucleic Acids Res 48(14):8074–8089 (2020). doi:10.1093/nar/gkaa549

83. X.H. Liang, S.T. Crooke. Depletion of key protein components of the RISC pathway impairs pre-ribosomal RNA processing. Nucleic Acids Res 39(11):4875–4889 (2011). doi:10.1093/nar/gkr076

84. A.A. Kelani, A. Bruch, F. Rivieccio, C. Visser, T. Krueger, D. Weaver, X. Pan, S. Schaeuble, G. Panagiotou, O. Kniemeyer, M.J. Bromley, P. Bowyer, A.E. Barber, A.A. Brakhage, M.G. Blango. Disruption of the Aspergillus fumigatus RNA interference machinery alters the conidial transcriptome. RNA 29(7):1033–1050 (2023). doi:10.1261/rna.079350.122

85. K. Spasov, L.I. Perdomo, E. Evakine, R.N. Nazar. RAC protein directs the complete removal of the 3’ external transcribed spacer by the Pac1 nuclease. Mol Cell 9(2):433–437 (2002). doi:10.1016/s1097-2765(02)00461-6

86. D.A. Bernstein, V.K. Vyas, D.E. Weinberg, I.A. Drinnenberg, D.P. Bartel, G.R. Fink. Candida albicans Dicer (CaDcr1) is required for efficient ribosomal and spliceosomal RNA maturation. Proc Natl Acad Sci USA 109(2):523–528 (2012). doi:10.1073/pnas.1118859109

87. A.K. Srivastava, D. Schlessinger. Processing pathway of Escherichia coli 16S precursor rRNA. Nucleic Acids Res 17(4):1649–1663 (1989). doi:10.1093/nar/17.4.1649

88. J.E. Burke, A.D. Longhurst, P. Natarajan, B. Rao, J. Liu, J. Sales-Lee, Y. Mortensen, J.J. Moresco, J.K. Diedrich, J.R. Yates, H.D. Madhani. A non-Dicer RNase III and four other novel factors required for RNAi-mediated transposon suppression in the human pathogenic yeast Cryptococcus neoformans. G3 (Bethesda) 9(7):2235–2244 (2019). doi:10.1534/g3.119.400330

89. L.C. Yeh, J.C. Lee. Structure analysis of the 5’ external transcribed spacer of the precursor ribosomal RNA from Saccharomyces cerevisiae. J Mol Biol 228(3):827–839 (1992). doi:10.1016/0022-2836(92)90867-j

90. J. Chen, L. Zhang, K. Ye. Functional regions in the 5’ external transcribed spacer of yeast pre-rRNA. RNA 26(7):866–877 (2020). doi:10.1261/rna.074807.120

91. H. Bourbon, B. Michot, N. Hassouna, J. Feliu, J.P. Bachellerie. Sequence and secondary structure of the 5’ external transcribed spacer of mouse pre-rRNA. DNA 7(3):171–191 (1988). doi:10.1089/dna.1988.7.181

92. M.H. Renalier, S. Mazan, N. Joseph, B. Michot, J.P. Bachellerie. Structure of the 5’-external transcribed spacer of the human ribosomal RNA gene. FEBS Lett 249(2):279–284 (1989). doi:10.1016/0014-5793(89)80641-6

93. M.R.E. Wong, K.H. Lim, E.X.Y. Hee, H. Chen, C.H. Kuick, S.J. Aw, K.T.E. Chang, N.S. Sulaiman, S.Y. Low, S. Hartono, A.N.T. Tran, S.H. Ahamed, C.M.J. Lam, S.Y. Soh, K.M. Hannan, R.D. Hannan, L.A. Coupland, A.H.P. Loh. Targeting mutant dicer tumorigenesis in pleuropulmonary blastoma via inhibition of RNA polymerase I. Transl Res 258:60–71 (2023). doi:10.1016/j.trsl.2023.03.001

94. B. Roche, B. Arcangioli, R. Martienssen. New roles for Dicer in the nucleolus and its relevance to cancer. Cell Cycle 16(18):1643–1653 (2017). doi:10.1080/15384101.2017.1361568

95. B. Bai, S. Yegnasubramanian, S.J. Wheelan, M. Laiho. RNA-seq of the nucleolus reveals abundant SNORD44-derived small RNAs. PLoS One 9(9):e107519 (2014). doi:10.1371/journal.pone.0107519

96. G.H. Wei, D. Dong, P. Zhang, M. Liu, Y. Wei, X. Wang, W. Xu, Q. Zhang, Y. Zhu, Q. Zhang, X. Yang, J. Zhu, L. Wang. Combined SNPs sequencing and allele specific proteomics capture reveal functional causality underpinning the 2p25 prostate cancer susceptibility locus. Res Sq [Preprint] (2024). doi:10.21203/rs.3.rs-3943095/v1

97. Y. Shima, K. Yamagata, Y. Kuroki, K. Sasaki, Y. Aikawa, I. Kitabayashi. Loss of NOL10 leads to impaired disease progression of NUP98::DDX10 leukemia. Leukemia 39(6):1368–1379 (2025). doi:10.1038/s41375-025-02607-5

98. S. Ben Hassine, B. Arcangioli. Tdp1 protects against oxidative DNA damage in non-dividing fission yeast. EMBO J 28(6):632–640 (2009). doi:10.1038/emboj.2009.9

99. M. Martin. Cutadapt removes adapter sequences from high-throughput sequencing reads. EMBnet J 17(1) (2011). doi:10.14806/ej.17.1.200

100. B. Langmead, S.L. Salzberg. Fast gapped-read alignment with Bowtie 2. Nat Methods 9(4):357–359 (2012). doi:10.1038/nmeth.1923

101. A.R. Quinlan, I.M. Hall. BEDTools: a flexible suite of utilities for comparing genomic features. Bioinformatics 26(6):841–842 (2010). doi:10.1093/bioinformatics/btq033

102. N.A. Joshi, J.N. Fass. Sickle: a sliding-window, adaptive, quality-based trimming tool for FastQ files. (2011) [Software]

103. H. Li, B. Handsaker, A. Wysoker, T. Fennell, J. Ruan, N. Homer, G. Marth, G. Abecasis, R. Durbin, 1000 Genome Project Data Processing Subgroup. The sequence alignment/map format and SAMtools. Bioinformatics 25(16):2078–2079 (2009). doi:10.1093/bioinformatics/btp352

104. E. Garrison, G. Marth. Haplotype-based variant detection from short-read sequencing. arXiv:1207:3907 (2012). doi:10.48550/arXiv.1207.3907

105. R. Sepsiova, I. Necasova, S. Willcox, K. Prochazkova, P. Gorilak, J. Nosek, C. Hofr, J.D. Griffith, L. Tomaska. Evolution of telomeres in Schizosaccharomyces pombe and its possible relationship to the diversification of telomere binding proteins. PLoS One 11(4):e0154225 (2016). doi:10.1371/journal.pone.0154225

106. Y. Oizumi, T. Kaji, S. Tashiro, Y. Takeshita, Y. Date, J. Kanoh, J. Complete sequences of Schizosaccharomyces pombe subtelomeres reveal multiple patterns of genome variation. Nat Commun 12(1):611 (2021). doi:10.1038/s41467-020-20595-1

107. A. Matsuda, Y. Chikashige, D.Q. Ding, C. Ohtsuki, C. Mori, H. Asakawa, H. Kimura, T. Haraguchi, Y. Hiraoka. Highly condensed chromatins are formed adjacent to subtelomeric and decondensed silent chromatin in fission yeast. Nat Commun 6:7753 (2015). doi:10.1038/ncomms8753

108. S. Tashiro, Y. Nishihara, K. Kugou, K. Ohta, J. Kanoh. Subtelomeres constitute a safeguard for gene expression and chromosome homeostasis. Nucleic Acids Res 45(18):10333–10349 (2017). doi:10.1093/nar/gkx780

109. M. Shimanuki, L. Uehara, T. Pluskal, T. Yoshida, A. Kokubu, Y. Kawasaki, M. Yanagida. Klf1, a C2H2 zinc finger-transcription factor, is required for cell wall maintenance during long-term quiescence in differentiated G0 phase. PLoS One 8(10):e78545 (2013). doi:10.1371/journal.pone.0078545

110. D.A. Bitton, F. Schubert, S. Dey, M. Okoniewski, G.C. Smith, S. Khadayte, V. Pancaldi, V. Wood, J. Bähler. AnGeLi: a tool for the analysis of gene lists from fission yeast. Front Genet 6:330 (2015). doi:10.3389/fgene.2015.00330

111. R. Lorenz, S.H. Bernhart, C. Höner zu Siederdissen, H. Tafer, C. Flamm, P.F. Stadler, I.L. Hofacker. ViennaRNA Package 2.0. Algorithms Mol Biol 6:26 (2011). doi:10.1186/1748-7188-6-26

112. M. Zuker, M. Mfold web server for nucleic acid folding and hybridization prediction. Nucleic Acids Res 31(13):3406–3415 (2003). doi:10.1093/nar/gkg595

113. R.V.A. Intine, thesis, University of Guelph, Canada. Structural features of the 5’ETS in Schizosaccharomyces pombe essential for ribosomal RNA maturation (1999).

114. J. Barandun, M. Hunziker, S. Klinge, S. Assembly and structure of the SSU processome – a nucleolar precursor of the small ribosomal subunit. Curr Opin Struct Biol 49:85–93 (2018). doi:10.1016/j.sbi.2018.01.008

115. M. Mann, P.R. Wright, R. Backofen, R. IntaRNA 2.0: enhanced and customizable prediction of RNA-RNA interactions. Nucleic Acids Res 45(W1):W435–W439 (2017). doi:10.1093/nar/gkx27

116. N.L. Bray, H. Pimentel, P. Melsted, L. Pachter. Near-optimal probabilistic RNA-seq quantification. Nat Biotechnol 34(5):525–527 (2016). doi:10.1038/nbt.3519

117. R. Schmieder, R. Edwards. Quality control and preprocessing of metagenomic datasets. Bioinformatics 27(6):863–864 (2011). doi:10.1093/bioinformatics/btr026

118. M.S. Scott, P.V. Troshin, G.J. Barton. NoD: a nucleolar localization sequence detector for eukaryotic and viral proteins. BMC Bioinformatics 12:317 (2011). doi:10.1186/1471-2105-12-317

119. H.S. Kim, B. Roche, S. Bhattacharjee, L. Todeschini, A.Y. Chang, C. Hammell, A. Verdel, R.A. Martienssen. Clr4SUV39H1 ubiquitination and non-coding RNA mediate transcriptional silencing of heterochromatin via Swi6 phase separation. Nat Commun15(1):9384 (2024). doi:10.1038/s41467-024-53417-9

120. K. Ekwall, T. Olsson, B.M. Turner, G. Cranston, R.C. Allshire. Transient inhibition of histone deacetylation alters the structural and functional imprint at fission yeast centromeres. Cell 91(7):1021–32 (1997). doi:10.1016/s0092-8674(00)80492-4

