## Supplemental information for "RiboCop surveils pre-rRNA processing by Dicer in cellular quiescence"

B. Roche, R.A. Martienssen

**The PDF file includes:**

Materials and Methods

Figs. S1 to S5

Tables S1 to S2

References

Materials and Methods

*S. pombe* strains and culture

A list of *S. pombe* strains used in this study is provided in Table S1. All cultures conditions were at 30°C. To induce G_0_, a *S. pombe* culture (prototrophic strain) is first grown in EMM minimal medium, then shifted to EMM-N at an initial OD=0.1, as described previously *(15)*. To measure cell viability, >100 individual cells (N=100 to 112) were micro-isolated using a Singer MSM400 micro-manipulator (Singer Instruments) and deposited unto a YES plate (rich medium), and the CFU (colony-forming unit) was counted after 3 days, as described before *(15,98)*. While this technique measures not only G_0_ viability but also ability to re-enter the cell cycle (mutants with a specific defect in G_0_-exit do not form colonies), in the case of Dicer, we have previously described that the majority of cells die in G_0_, as evidenced by the large quantity of apoptotic cells without DNA in *dcr1*Δ cultures after 2 weeks of G_0_ and the inability of non-colony forming cells to initiate S-phase when plated on rich medium *(15)*. Common viability dyes (phloxin B, propidium iodide, trypan blue) do not stain G_0_ cells efficiently and therefore cannot be used to reliably assay G_0_ viability. We typically use strains of mating-type h- because of its stability over h+, but both mating-types display comparable viabilities in EMM-N. Deletion mutants were generated by transformation using the EZ Zymo kit (Zymo Research), using primers provided in Table S2, and combination of mutants were generated by crosses on ME plates. To assay the viability of Enp2 mutants, a diploid strain was obtained following mating of compatible *ade6*-M210 and *ade6*-M216, and used for transformations; the resulting heterozygote was isolated and sporulated, followed by tetrad analysis using the Singer MSM400 micro-manipulator (Singer Instruments).

Improved parallel analysis of RNA 5’ ends and sequencing (iPARE-seq)

For iPARE-seq libraries, total RNA was isolated from 100ml of G_0_ cells (EMM-N medium) using the ZR Fungal/Bacterial RNA Miniprep kit (Zymo Research) according to manufacturer’s instruction, keeping small RNAs (2x volumes ethanol used), including an in-column DNase I digestion step 20 minutes at room temperature, and eluted in 21μl H_2_O. 1μl of RNA was used for quantification by fluorometry using a QuBit 2.0 fluorometer (Thermo Fischer Scientific) with the RNA Broad Range assay kit (Thermo Fischer Scientific). 1μg of total RNA was ligated to the 5’ PARE adapter (100pmol) in 10% DMSO, 1mM ATP, 1X T4 RNA ligase 1 buffer (NEB), 25% PEG8000 with 1μl (40U) of RNase OUT (Thermo Fischer Scientific) and 1μl (30U) of T4 RNA ligase 1 (NEB) in a total volume of 100μl. The ligation was performed for 2 hours at 25°C followed by an overnight incubation (~18 hours) at 16°C, then purified using RNA Clean XP beads (Beckman Coulter) according to manufacturer’s instructions, eluting in 18μl H_2_O. Ligated RNA was chemically fragmented using the Magnesium RNA fragmentation kit (New England Biolabs) to reduce the size of the captured fragments to ≤200nt, by adding 2μl of RNA Fragmentation Buffer, incubating 5 minutes at 94°C followed by a transfer to ice and the addition of 2μl of RNA Stop solution. The fragmented RNA was then purified using the RNA Clean & Concentrator-5 kit (Zymo Research) following manufacturer’s instruction, eluting in 11μl H_2_O. 10μl of RNA were used in the reverse-transcription reaction as follows: first, 1μl of dNTP (10mM) (Invitrogen) and 2μl of Random primer mix (New England Biolabs; contains both N_6_ hexamers and dT_23_VN anchored primers) were added, the reaction mixed and incubated 5 minutes at 65°C, then put on ice for 1 minute. Afterwards, 4μl of SuperScript IV buffer (5X) (Thermo Fischer Scientific), 1μl of DTT (100mM), 1μl of RNaseOUT (Thermo Fischer Scientific) and 1μl of SuperScript IV (200U) (Thermo Fischer Scientific) were added, and the reaction was incubated 10 minutes at 23°C followed by 10 minutes at 50°C. Finally, 80μl of TE was added. To perform target indirect capture, we first washed 100μl streptavidin magnetic beads (Dynabeads MyOne Streptavidin T1, Thermo Fischer Scientific) by adding 500μl of BW buffer (5mM Tris-HCl pH=7.5, 500μM EDTA, 1M NaCl), incubating at room temperature on a rotating platform for 5’, followed by magnet separation for 2’ and removal of the wash buffer. 2 more washes were performed with 500μl BW buffer each, followed by 2 washes with 500μl A buffer (100mM NaOH, 50mM NaCl), followed by 4 washes with B buffer (100mM NaCl), then the beads were finally resuspended in 100μl of 2xBW buffer. 100μl of the RT reaction was added and mixed well, followed by room temperature incubation for 15’ with rotation. The beads were washed 3 times with 500μl BW buffer, followed by 3 washes with 700μl 1xSSC buffer. Finally, the beads were resuspended in 50μl of 1xSSC buffer, incubated at 95°C for 5’, magnet-separated for 2’ and the 50μl containing the captured cDNA molecules were transferred to a new tube. Second-strand synthesis was performed using Klenow fragment (5U) (NEB) with 100μM dNTPs and 1μM of the PARss adapter primer 5’-NNNNTCTAGAATGCATGGGCCCTCCAAG-3’ for 1h at 37°C, followed by heat inactivation at 75°C for 20 minutes. The reaction was further purified using AMPure XP SPRI magnetic beads (Beckman Coulter) (ratio 1:1) following manufacturer’s instruction, resuspended in 51μl EB. 1μl was used for quantification using the QuBit 2.0 fluorometer (Thermo Fischer Scientific) and the remaining 50μl were used for library generation using the NEB Ultra DNA library kit (NEB). The libraries were sequenced on a NextSeq (Illumina), paired-end run type (151nt + barcode), except for one replicate of *wt* and one replicate of *dcr1*Δ which were sequenced on a MiSeq, paired-end run type (301nt + barcode). Including or excluding these replicates did not affect our conclusions. Because the iPARE adapter is directional, using the non-directional DNA-seq kit does not result in a loss of information, but the adapter must be searched for either in 5’ (forward orientation in R1) or in 3’ (forward orientation in R2). To sort these reads, we used CUTADAPT to detect the adapter in R1 (--no-trim parameter) *(99)*. R1 adapter reads were then trimmed for the 3’ adapter if present, using CUTADAPT (5’-AGATCGGAAGAGCACACGTCTGAACTCCAGTC-3’, minimum overlap 8nt). For each read the 4 N bases were recorded, analyzed by a custom perl script to assay bias, followed by trimming the 5’ iPARE adapter with CUTADAPT (5’-NNNNTCTAGAATGCATGGGCCCTCCAAG-3’), then trimming potential poly-A tails in 3’ (option -a A{60}), then trimming potential NextSeq ‘dark cycles’ appearing as poly-G tails in 3’ and finally keeping only reads ≥ 20nt (option -a G{60} -m 20). These reads were further quality-filtered using FASTXTOOLKIT keeping a minimum quality of 20 over 70% of bases. R2 adapter reads were detected similarly (--no-trim parameter, -G adapter option) and processed in an identical pipeline. Quality-filtered reads were mapped as single-end reads using BOWTIE2 *(100)*, and the genome coverage of 5’ positions was calculated using BEDTOOLS *(101)* with RPM normalization. While in the presented data all mapped reads were kept, the optional inclusion of a deduplication step based on the N4 using a custom perl script did not affect our results and conclusions. Composition analysis of these 4 N bases did not show bias. To detect peaks along the mature rRNA regions, we detected local maxima at these intervals (excluding the mature rRNA 5’ ends) in two steps: (i) first, positions with a minimum coverage (RPM≥400) were selected; (ii) then, only positions with higher coverage than both the previous (-1nt) and the next (+1nt) nucleotide position, and for which the highest neighboring nucleotide position had no more than 80% of the RPM signal of that maximum. Each peak was then further manually confirmed. The same approach was used to detect peaks at the pre-rRNA regions, except that the RPM threshold was lowered to ≥1 to account for the significant difference in RNA level of pre-rRNA vs. rRNA. A similar approach was used to detect peaks over the rest of the genome, with a RPM threshold of ≥2 where local maxima were detected in windows of 41nt (±20nt). Peaks were detected independently for wild-type, *dcr1*Δ and *dcr1-5* replicates, and combined (to allow detection of peaks that would be completely Dicer-dependent or Dicer-induced). Pre-rRNA peaks were further checked manually.

Chromatin immunoprecipitation.

Chromatin immunoprecipitation (ChIP) was performed as previously described *(15)* as a modification of the Zymo-Spin ChIP kit protocol (Zymo Research). The cross-linking reaction was done at room temperature with shaking for 10 mins for H3K9me2 using the ab1220 antibody (Abcam), for 30 mins for FLAG-tagged RNA pol I (*nuc1*-(Gly)_6_-3xFLAG and *rpa2*-(Gly)_6_-3xFLAG) using the M2-anti-FLAG antibody (Sigma). The purified ChIP DNA was then used for qPCR, in which case the input samples were diluted 1:100, or for next-generation sequencing, which was performed according to the recommendations of the NEB Ultra II DNA-seq kit (New England Biolabs). Libraries were pooled in equimolar amounts and sequenced on a MiSeq platform (Illumina). FASTQ files resulting from sequencing were adapter-trimmed using CUTADAPT *(99)*, quality-filtered using SICKLE *(102)*, mapped using BOWTIE2 *(100)*, deduplicated using SAMTOOLS *(103)* and normalized genome coverage was computed using BEDTOOLS *(101)*.

For ChIP-exo, 100ml of culture in EMM-N medium (2 days of G_0_) was fixed by addition of formaldehyde to a final concentration of 1% (+2.7ml 37%) for 30 minutes at room temperature with frequent mixing. The crosslinking reaction was quenched by the addition of 5ml of glycine (2.5M) for 10 minutes followed by centrifugation for 2 minutes at 3000rpm. The cell pellet was washed twice with PBS. Cell lysis and immunoprecipitation were performed identically to the ChIP protocol. The immunoprecipitation was performed overnight with either 2μl of mouse monoclonal anti-FLAG M2 antibody (Sigma Aldrich) or a rabbit anti-IgG control (Cell Signaling Technology), in a rotator. 15μl of protein A beads (Zymo Research) were added to the sample, incubated for 1h at 4°C and washed according to the kit (Zymo Research). After the last wash, the pellet was resuspended in the end-repair on-bead reaction mix (total volume 50μl): 37μl H2O, 5μl CutSmart 10X buffer (NEB), 2μl PNK (10U/μl) (NEB), 2μl T4 DNA polymerase (3U/μl), 1μl Klenow polymerase (5U/μl), 3μl of dNTPs (1mM each) and 0.5μl ATP 10mM. The end-repair reaction was incubated at 37°C for 1h, followed by an inactivation step 5 minutes at 70°C. The following reagents were then added to the mix for the on-bead exonuclease reaction: 41μl H2O, 5μl CutSmart 10X buffer (NEB) and 4μl λ-exonuclease (5U/μl) (NEB), and incubated at 37°C for 1h. The reaction was inactivated 15 minutes at 75°C, after which 400μl of 1X Chromatin Elution Buffer (Zymo Research) and 20μl of NaCl (6M) were added and kept 15 minutes at 75°C. The magnetic beads were pelleted by a short centrifugation (30 seconds at 10,000 rpm) and the supernatant transferred to new tubes. Reverse-crosslinking was then performed by incubating the samples 4h30 at 65°C, followed by 10 minutes at 95°C and 10 minutes at room temperature. Following reverse crosslinking, chromatin was column-purified using the Zymo ChIP Clean & Concentrate kit (Zymo Research), with the final elution step using 2x10μl water. The purified DNA was then sequenced using a single-strand DNA-seq protocol, using the Accel-NGS 1S Plus DNA library kit (Swift Biosciences) following manufacturer’s instructions. The SPRI magnetic bead (Agencourt AMPure XP beads, Beckman-Coulter) clean-up ratios were adjusted to optimize the cut-off for small fragments in the following manner: the 1^st^ bead clean-up (post-adaptase reaction step #13) was adjusted to a ratio of 1.6X (87μl sample + 139μl magnetic beads) and the 2^nd^ bead clean-up (post 5’-ligation, step #17) was adjusted to a ratio of 1.4X (40μl sample + 56μl magnetic beads). The libraries were made with distinct barcodes from the Accel-NGS 1S Plus DNA library kit (Swift Biosciences), followed by qPCR quantification using the Kapa Library Quantification kit for Illumina (Kapa Biosystems), after which equimolar amounts were pooled for sequencing using an Illumina MiSeq, run-type single-read 50nt with barcode. Following sequencing, FASTQ files of ChIP-exo reads were adapter-trimmed using CUTADAPT *(99)* in two steps: first, the 3’ adapter is removed using the following parameters (-a AGATCGGAAGAGCACACGTCTGAAC -m 20 -n 6 --trim-n -O 6); then, the 3’ tail incorporated by adaptase (cf. Swift Biosciences documentation for the Accel-NGS 1S kit) was cleaved using CUTADAPT (parameter -u -10). The trimmed reads were then mapped to the *S. pombe* genome using BOWTIE2 *(100)*, sorted using SAMTOOLS *(103)* and the genomic coverage of the 5’ positions (corresponding to the protein footprint) was calculated using BEDTOOLS *(101)*. Read counts were normalized to RPM values and visualized using custom R scripts.

Characterization and fine-mapping of the *deltel1R* interval

The *deltel1R* was recovered as a spontaneous suppressor in a microevolution screen alternating a *dcr1*Δ culture between growth and quiescence *(15)*. Genomic DNA from the suppressed strain was prepared using the Genomic Tip 25/g (Qiagen), and libraries were prepared using the Illumina PCR-free TruSeq kit (Illumina), sequenced on a MiSeq (Illumina) (run-type: paired-end 151nt), and the analysis performed as described before *(15)*. Briefly, paired reads were 3’-trimmed using SICKLE *(102)*, quality-filtered using FASTXTOOLKIT, mapped to the *S. pombe* genome using BOWTIE2 *(100)* then sorted using SAMTOOLS *(103)*. We did not find SNPs in this strain compared to the initial wild-type and/or in the initial *dcr1*Δ strain used to start the suppressor screen, using FREEBAYES *(104)*. By plotting the DNA-seq coverage in 1kb windows over the whole genome to look for copy number variations (CNVs), we identified a large deletion proximal to the right telomere of chromosome 1 (fig. S3A). To narrow the left-point of the deletion, we repeated the plot using 5nt windows. To identify split-reads, we extracted all reads that were unmapped by BOWTIE2 (each read in a pair now being treated independently). All reads ≥60nt long were then cut to keep 25nt at the 5’ and 25nt at the 3’ end, these small reads were mapped independently using BOWTIE2 (in single-end mode), and re-assembled in order to search for reads in which one end maps at the narrowed region (between chr1:5439000 and chr1:5440000) while the other end does not map; this approach uncovered 7 ‘split-reads’. Multiple sequence alignment between these reads and the chr1:5439000-5440000 genomic sequence of *S. pombe* shows that all 7 reads perfectly map to *S. pombe* up to chr1:5439552, and are then followed by an identical different sequence which consists of up to eight and a half repetitions of the *S. pombe* heterogeneous telomeric repeat (G_2-9_TTAC[A][C])_n_ *(105)*. Furthermore, the sequence of the ‘mate’ read for each of these split-reads was a reverse-complemented telomeric repeat, as expected. The location of the deletion was confirmed by PCR using primers del-A and del-B on either side of the breakpoint, which only amplified in wild-type cells and the original *dcr1*Δ strain that was used as a starting point of the screen, but not in the recovered *dcr1*Δ *deltel1R* suppressor nor in a back-crossed *deltel1R* strain (fig. S3B). Inversely, PCR amplification using del-A and the telomeric primer del-T resulted in amplification only in *deltel1R* strains (fig. S3B). Furthermore, the reduction of copy number of ST genes by loss of ST1R was confirmed by qPCR on *SPAC750.02c* and *ftm4* as compared to *act1*, as well as to *clr5* and *SPAC29B12.13*, which are genes proximal to *deltel1R* (data not shown). The *deltel1R* deletion is ~139kb long based on the current *S. pombe* assembly, with an estimated additional ~19kb missing from the assembly as determined by recent complete subtelomeric sequencing *(106)*, for a total deletion size of ~158kb. Overall, this region encompasses 46 protein-coding genes and an estimated 34 ncRNAs, and reducing the copy number of a repeated block of 7 sub-telomeric genes normally found at both ends of chromosome 1 and 2. The deletion starts 3’ of chr1:5439552 and spans the rest of the subtelomeric region up to the telomeric repeats, as evidenced by the recovery of ‘split-reads’, spanning the deletion site and eight and a half telomeric G_2-9_TTAC[A][C] repeats *(105)*. As the deletion encompasses the ST1R and SH1R highly-condensed sub-telomeric regions *(107,108)*, we assayed whether this structural variation leads to meiotic defects. We compared the spore viability in *wt* x *deltel1R* crosses, where the structural arrangement differs, versus *wt* x *wt* and *deltel1R* x *deltel1R* crosses, performing tetrad analysis using a Singer MSM400 micro-manipulator (Singer Instruments) and did not observe any significant difference in spore viability and crossing efficiency (data not shown). Only full tetrads (4 spores) were taken into account for this analysis; a minimum of 68 tetrads were isolated. The inclusion of tetrads with less than 4 spores does not affect these results as only two were found amongst 215 full tetrads analyzed (0.9%). This result is in agreement with a previous study that found that deletion of all ST repeats does not cause meiotic defects *(108)*. The most likely scenario is therefore that the suppression by *deltel1R* is conferred through the deletion of a gene within the region. The only *a priori* candidate locus within *deltel1R* was the transcription factor Klf1 (fig. S3A), which has been previously described to be important for long-term G_0_ viability *(109)*, but we assayed the phenotype of *klf1*Δ and *dcr1*Δ*klf1*Δ mutants in G_0_, and did not find any suppression effect (fig. S3C). We noticed that the *deltel1R* region contains one Dicer-dependent ncRNA, NC30 (fig. S3DE), which was also a candidate for ncRNAs displaying complementarity to the pre-rRNA spacers (fig. S3I), making it a strong candidate. We therefore constructed deletion mutants of this ncRNA, as well as two control ncRNAs from *deltel1R*, NC280 and NC281, and assayed the viability of the *dcr1*Δ*NC30*Δ, *dcr1*Δ*NC280*Δ and *dcr1*Δ*NC281*Δ double-mutants. We found that *dcr1*Δ*NC30*Δ displayed a partial suppression of quiescence maintenance defects (Fig 2B). In contrast, the double-mutants *dcr1*Δ*NC280*Δ and *dcr1*Δ*NC281*Δ did not display any suppression, nor did deletion of their neighboring genes *dcr1*Δ*yhb1*Δ and *dcr1*Δ*SPAC869.03c*Δ (fig. S3J). To assay if the suppression observed in *dcr1*Δ*NC30*Δ is due to a *cis*-effect on neighboring gene expression (that is, transcriptional interference or promoter destabilization), we constructed the deletion mutants *dcr1*Δ*gto1*Δ and *dcr1*Δ*SPAC922.03*Δ, and neither displayed suppression of the *dcr1*Δ G_0_ phenotypes (fig. S3J).

To characterize the RiboCop promoter (fig. S3H), we searched for all HomolD boxes present on the *S. pombe* genome (excluding the rDNA promoter) using a custom perl script, and found 244 boxes. Their repartition between chromosomes was not significantly different than random (χ^2^ test, p=0.94). Protein-coding genes located within 100nt downstream of a HomolD box (82/244) were, as expected, highly enriched for the GO term ‘ribosome biogenesis’ (p-value<7×10^-12^) and corresponded to genes regulated by the ‘ribosomal protein module HomolD and E’ (p-value<2.2×10^-51^), which are typically highly-expressed (p-value<3.5×10^-47^), repressed by environmental stresses (p-value<4.6×10^-30^), caffeine and rapamycin (p-value<1×10^-16^; enrichments were calculated using AnGeLi *(110)*). We found 7 snoRNAs (snR41, snoR02, snoR56, snoZ15, snR42, sno20, and the U3A snoRNA snu3) and 28 ncRNAs located within 200nt downstream of a HomolD box, 8 of which share a bidirectional promoter with protein-coding genes or snoRNAs involved in ribosomal biogenesis. The overlap between these 28 ncRNAs and the 91 ncRNAs up-regulated in *dcr1*Δ consisted of only two ncRNAs, NC30 and NC1080. The overlap is not statistically significant (Fisher’s exact test, p=0.36), which argues that Dicer does not have a general role in regulation of HomolD ncRNAs, but rather specifically of NC30 and NC1080.

RNA secondary structure predictions and candidate lncRNA:pre-rRNA interactions

The RNA secondary structure for NC30/RiboCop was predicted using RNAFOLD *(111)* and MFOLD *(112)*. pre-rRNA structures in the 5’ETS and 3’ETS (Fig. 1EF) was determined by manual comparison with *S. cerevisiae* 5’ETS hairpins, successive rounds of RNAFOLD prediction for each 5’ETS hairpin independently and in pairs, and comparison with previous work *(35,113)*. The 3’ETS structure was determined by RNAFOLD and comparison with a previously proposed structure *(28)*. While a 5’ETS processosome structure was recently published from *S. cerevisiae (114)*, the significant sequence divergences in spacer regions did not allow us to gain information on the tertiary structure for the *S. pombe* 5’ETS.

To screen for potential lncRNA:pre-rRNA interactions, we selected complete sequences from all Dicer-dependent ncRNAs identified from our RNA-seq experiment, and split the full 5’ETS pre-rRNA sequence (chr3:2439693-2440993) in 50nt windows (sliding by 20nt). A custom perl script performed the following operation for every combination: first, the ncRNA was reverse-complemented and a perfect match with the rRNA window was searched (minimum *k*-mer size: 5nt). Every match was extended as possible to determine the longest *k*-mer size with a perfect match (maximum *k*-mer size: 50nt). Then, the ncRNA sequence was randomly shuffled (by assembling random subsets of the ncRNA using random 5nt windows, concatenated until the ncRNA size is reached) and the same procedure is applied to this ‘randomized’ ncRNA to find its longest perfectly matched *k*-mer. The randomization procedure is repeated for a total of 100 bootstraps and the average ‘randomized’ *k*-mer value is obtained, as well as its standard deviation (typically ±1nt). In figure S3I, the heatmap color represents the difference between the longest *k*-mer for the original ncRNA vs. the average longest *k*-mer of the randomized ncRNA, for every position, i.e. highlights positions where the ncRNA:pre-rRNA reverse-complementarity appears higher than by chance. The best predicted interactions were further manually checked using INTARNA *(115)*. Within *deltel1R*, NC30/RiboCop was the ncRNA with the best candidate interaction site. The very low expression of NC30/RiboCop, and very high expression of rRNA, did not allow us to unambiguously confirm this interaction *in vivo* using RNA:RNA crosslinking.

RNA-seq and small RNA-seq

Total RNA was purified from 100ml cell cultures 48h after G_0_ induction (EMM-N medium), using the ZR Fungal/Bacterial RNA Miniprep kit (Zymo Research), following manufacturer’s instructions—including the in-column DNase I digestion step (20 minutes at room temperature). Ribosomal RNA fragments were depleted using the Ribo-Zero rRNA removal kit (human/mouse/rat) (Illumina) and libraries made using the ScriptSeq v2 non-directional RNA-seq kit (Epicenter) following manufacturer instructions. Barcoded RNA-seq libraries were quantified using the Kapa Quantification Kit (Kapa Biosystems), pooled at equimolar amounts and sequenced on an Illumina NextSeq, using a high through-put run-type of single-read 76nt + barcode. The library 3’ adapter (5’-AGATCGGAAGAGCACAGTCTG-3’) was clipped from the reads using CUTADAPT, version 2.0 *(99)*, using options -O 6 -n 2 -j 4 --next-seq-trim 20 --trim-n, followed by trimming of 3’-terminal poly(A) tails using CUTADAPT with options -a A{60} -O 30 -j 4, then followed by trimming of 3’-terminal poly(G) ‘dark cycles’ resulting from the two-color NextSeq chemistry using CUTADAPT with options -a G{60} -O 30 –j 4. Reads were quality-filtered using FASTXTOOLKIT version 0.0.13.2 (<http://hannonlab.cshl.edu/fastx_toolkit/)>. Reads were aligned to the *S. pombe* transcriptome (excluding 18S, 5.8S and 28S rRNA sequences) using KALLISTO version 0.43.1 *(116)*. RNA-seq libraries were constructed in triplicate for wild-type and *dcr1*Δ G_0_ cells.

To test for enrichment of Gene Ontology (GO) terms in Dicer-dependent transcripts, we used the online tool AnGeLi from the Bähler lab (<http://bahlerweb.cs.ucl.ac.uk/cgi-bin/GLA/GLA_input>) *(110)*. Enrichment was calculated independently for the upregulated and downregulated gene list, including either all genes or all protein-coding genes, against each available category and GO database (biological process, cellular component, molecular function). No significant enrichment was found for genes up-regulated in *wt* vs. *dcr1*Δ. For genes upregulated in *dcr1*Δ, in addition to the clear enrichment in non-coding RNAs which was also detected by this approach (p-value<1.2×10^-24^), there was a marked enrichment for intron-less genes (91%) of shorter length, with low expression level in both quiescent and dividing cells, and an enrichment for genes corresponding to the following datasets: “induced in Dbr1 deletion”, “core oxidative stress response”, “Pap1 but not Prr1-dependent genes”. This result suggests that in addition to ncRNAs, the stress response is active in *dcr1*Δ G_0_ cells, an expected result considering the strong viability loss and DNA damage accumulation observed in *dcr1*Δ mutants.

For small RNA-seq libraries, total RNA was purified from 100 ml of G_0_ cultures (48h EMM-N) using the ZR Fungal/Bacterial RNA Miniprep kit (Zymo Research), and small RNA-seq libraries were generated using the NEXTFLEX small RNA-seq kit v3 following manufacturer’s instructions (Bioo Scientific). Barcoded libraries were quantified using the Kapa Quantification Kit (Kapa Biosystems), pooled at equimolar amounts and sequenced on an Illumina MiSeq (run-type: single-end 50bp plus barcode). The 3’ adapter (5’-TGGAATTCTGGGTGCCAAGG-3’) was clipped using fastx_clipper from FASTXTOOLKIT and keeping a minimum size of 20nt (corresponding to 12nt of sequence + 4 N nucleotides in both 5’ and 3’). The composition of the 4 N nucleotides in 5’ and 3’ was assayed using a custom perl script, and showed a preference towards C at the last base of the 3’ N4 adapter (~46%), highlighting the importance of using N4 adapters to reduce ligation bias. The library was then deduplicated using PRINSEQ-lite version 0.20.4 *(117)* using the option -derep 1. The 4 N nucleotides were then removed from the 3’ end using CUTADAPT (parameters -u -4) *(99)*, followed by another round to remove the 5’ end 4 N nucleotides (parameters -u 4). The 5’ and 3’ bias of the clipped reads was then assayed using a custom perl script, and showed a slight bias towards A/U in 5’ (~60%) and no bias in 3’. The reads were then quality-filtered using FASTXTOOLKIT. Reads were mapped to the genome using BOWTIE2 version 2.3.3 (parameters -k 1 --no-unal) *(100)*, sorted using SAMTOOLS version 1.5 *(103)*, and the coverage over the genome was then calculated using BEDTOOLS version 2.29 *(101)* with RPM normalization with and without taking into account rDNA regions. For comparisons to RNA-seq values over determined regions (Dicer-dependent non-coding RNAs in fig. S3FG), strand-specific coverage was calculated using BEDTOOLS, and then sRNA-seq RPM values were normalized to feature length (in kb) to obtain RPKM values. For each region, we also calculated a ‘sense/antisense-ratio’ score as follows:

$$ratio= \frac{min(RPKMsense, RPKMantisense)}{max(RPKMsense, RPKMantisense)}$$

with the ratio being defined as 0 if both RPKM values are 0. As a result, this ratio is contained in the interval [0,1] with 0 corresponding to a region producing small RNAs only from one strand and 1 corresponding to an equal amount of small RNAs produced from both strands. Highly-transcribed genes generate sense degradation products, with a strong small RNA-seq signal but a sense/antisense ratio approaching 0, while canonical RNAi targets such as centromeres have high small RNA levels, very low expression levels (RNA), and a sense/antisense ratio approaching 1. This ratio measure is only accurate when there are enough reads in the considered region. As expected, there is a correlation between RNA level and small RNA levels in wild-type G_0_ cells (Spearman’s ρ=0.50), but an inverse correlation between small RNA levels and the sense/antisense-ratio (Spearman’s ρ=–0.38). The genes showing both a high level of Dicer-dependent small RNAs and a high sense/antisense ratio correspond in majority to known RNAi targets i.e. centromeric ncRNAs, centromeric boundary ncRNAs and the telomere-linked *tlh1*, *tlh2* and *SPAC212.06c* genes (fig. S1A).

We separated Dicer-dependent genes into four classes: (i) protein-coding genes, (ii) antisense non-coding RNAs (which were defined as non-coding RNAs located on the opposite strand of a protein-coding gene, with a minimum reciprocal overlap of 10%), (iii) centromeric non-coding RNAs, and (iv) other non-coding RNAs. Ranking genes based on the difference between their RNA and small RNA signal in wild-type G_0_ cells show that centromeric ncRNAs and the telomere-linked genes *tlh1* and *SPAC212.06c* are the main loci producing siRNAs. Overall, only centromeric ncRNAs had a significantly higher sRNA level difference between *wt* and *dcr1*Δ (i.e. ‘sRNA effect size’) compared to non-Dicer-dependent genes (one-tailed Mann-Whitney test, p<1.3×10^-6^), as well as a higher sense/antisense ratio (median ratio=0.63 vs 0.17 for all genes, one-tailed Mann-Whitney test p<7×10^-5^). Thus, the majority of small RNAs in other Dicer-dependent genes appear to represent degradation products rather than *bone fide* Dicer-dependent siRNAs, although we cannot discard the possibility that small RNAs are produced at very low levels at some of these loci.

NC30/RiboCop trans-expression experiments

For experiments to test whether RiboCop is able to exert its toxic effect in *trans*, we re-inserted the intergenic sequence, containing NC30/RiboCop and its promoter and terminator sequences, between the 5’UTR of *gto1* and the 5’UTR of *SPAC922.03* (UTR included but excluding any of their coding sequence), consisting of the chr1:5474549-5474579 interval, at different distinct genomic locations of chr1: (i) between *arg3* and *arg11* (ii) between *leu2* and *prl12*. To this effect, the *deltel1R* strain was transformed with the transgene using the EZ Zymo kit (Zymo Research), and crossed to a *deltel1R* *dcr1*Δ strain to obtain the triple-mutant. The loss of viability in quiescence in the triple-mutant *deltel1R* *dcr1*Δ *arg3::RiboCop* was not caused by a disruption of the *arg3* or *arg11* genes in *cis*, as our strains were not arginine auxotrophs but could grow in minimal medium (EMM). Similarly, we did not observe disruption of *leu2* in *leu2::RiboCop* in our strains, which would manifest as leucine auxotrophy. Furthermore, the *deltel1R arg3::RiboCop* and *deltel1R leu2::RiboCop* strains did not display a G_0_ phenotype, in accordance to the *trans*-effect being only triggered in the absence of *dcr1*.

Purification of NC30/RiboCop-associated proteins

To purify candidate proteins associated with the RiboCop riboprotein complex, given the very low expression level of the RiboCop lncRNA that precluded obtaining enough material from purifications of the endogenous complex, we opted for a ‘fishing-out’ strategy by combining *in vitro* transcribed RiboCop with protein extracts from *dcr1*Δ cells. First, a PCR product was amplified from BR00 genomic DNA to obtain the NC30/RiboCop sequence with a T7 promoter, and purified using the DNA Clean & Concentrate kit (Zymo Research). 800ng of purified PCR product was in vitro transcribed by T7 polymerase (New England Biolabs) following manufacturer’s instruction for 2h at 37°C (final 1X reaction buffer, 5mM DTT, 500μM each NTP, 1U/μl RNase OUT ribonuclease inhibitor (Thermo Fisher Scientific)). The reaction was stopped by denaturation 2 mins at 80°C, DNase I-treated 30 mins at 30°C, and purified using the Direct-zol Miniprep kit (Zymo Research). The purified RNA was treated with CIP to dephosphorylate the 5’P, with 1μl RNase OUT ribonuclease inhibitor (Thermo Fisher Scientific), followed by purification using the Direct-zol Miniprep kit (Zymo Research). T4 RNA ligase 1 was then used with pCp-biotin (Jena Biosciences) to 3’-biotinylate the RNA, for 2h 25°C followed by overnight incubation at 16°C (final: 50mM Tris HCl pH7.8, 10mM MgCl2, 1mM DTT, 1mM ATP, 10% DMSO, 1μM pCp-biotin, 10U T4 RNA ligase 1), and purified using the Direct-zol Miniprep kit (Zymo Research). Purified NC30 was folded *in vitro* in 3.1 buffer (New England Biolabs) by incubating 2mins 90°C, 30 mins at 37°C and 30 mins at room temperature. To obtain the protein extract, *dcr1*Δ cells (1L of late log-phase culture in YES) were pelleted and resuspended in lysis buffer (mTEG: 40mM Tris HCl pH 7.6, 1mM EDTA, 300mM NaCl, 10% glycerol, 0.1% NP-40) containing protease inhibitors (1 protease inhibitor tablet per 5ml, 1mM PMSF, 2μM leupeptin, 2μM pepstatin, 40μg/ml antipaine, 1μg/ml aprotinin). Note that as G_0_ cells display high protease activity, this combination of protease inhibitors was necessary to limit degradation in the lysates; the concentration was optimized by assaying different buffer compositions using Western blots with the Nuc1-(Gly)_6_-3xFLAG background, as Nuc1 is a large protein (189 kDa) sensitive to degradation (data not shown). Cells were lysed using an equivalent amount of zirconia beads by alternating vortexing for 1min, followed by 1min on ice, repeated a total of 24 times, and the supernatant was transferred to new tubes and cleared by centrifugation at 12,000g for 5 minutes. 10mg of biotinylated, folded RiboCop RNA was incubated with 1.5ml of cell lysate at room temperature for 2h at 4°C in a rotator, in the presence of 10μl RNase OUT ribonuclease inhibitor (Thermo Fisher Scientific). Pull-downs were performed using 50μl of streptavidin beads (Dynabeads MyOne Streptavidin T1, Thermo Fisher Scientific), washed and prepared following manufacturer’s instructions (for RNA work), resuspended in mTEG buffer. The beads were washed extensively with mTEG buffer (6 times: 5 minutes at 4°C on a rotator, 5 minutes on a magnetic rack for separation). Elution was performed in 10ul of mTEG buffer containing 1M NaCl to limit dissociation of free streptavidin from the beads. An aliquot of the cell lysate, and flow-through for each incubation step were kept, as well as the final eluate, resuspended in Laemmli buffer (Biorad), and run on SDS-PAGE 4-20% gels (Biorad) followed by silver staining following manufacturer’s instructions (Biorad) or, in another iteration, by Coommassie staining. Visible bands were cut with a scalpel and sent to the Mass Spectrometry facility at the University of Nebraska-Lincoln for identification by LC-MS. The number of peptides was compared to the protein abundance (PomBase) to determine relative enrichment in the pull-down. To confirm the identity of the proteins, the entire workflow was repeated in *dcr1*Δ protein extracts with FLAG-tagged genetic backgrounds (*rnh1*-(Gly)_6_-3xFLAG, *pabp*-(Gly)_6_-3xFLAG, and *enp2*-(Gly)_6_-3xFLAG, cf. Table S1 for strains used in this study), with the gel used for a Western blot using the anti-FLAG M2 monoclonal antibody (F3165, Sigma). In the case of Rnh1, in order to ascertain that the interaction was not an artefact resulting from the association of *in vitro* transcribed RiboCop RNA with remaining DNA in the lysate, the experiment was repeated in the *dcr1*Δ*RiboCop*Δ *rnh1*-(Gly)_6_-3xFLAG background, in which the excess RiboCop RNA cannot form a RNA:DNA hybrid with its endogenous locus (Fig 4D).

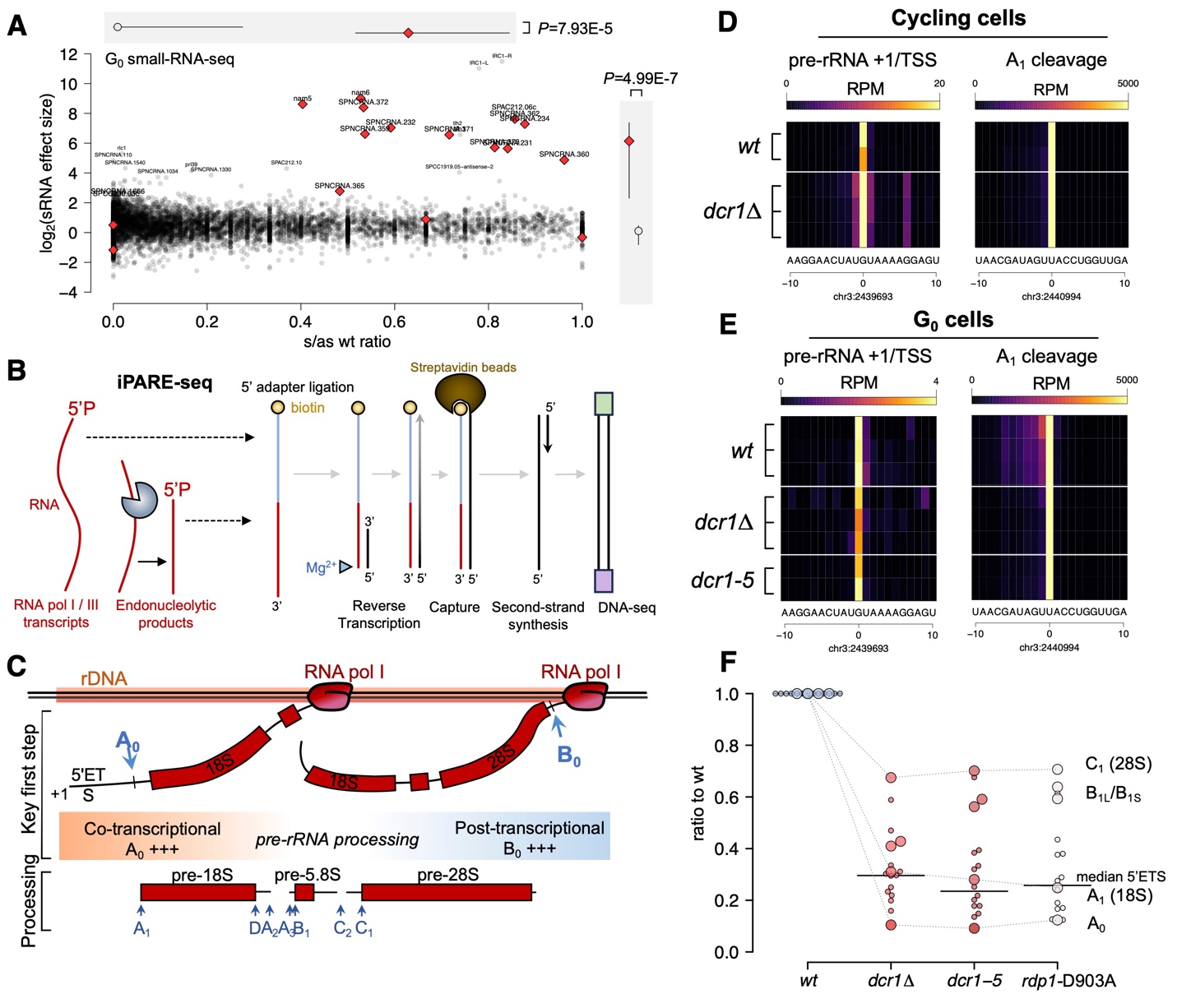

Fig. S1. Single-nucleotide detection of pre-rRNA cleavage sites by iPARE-seq.

(**A**) small RNA signatures are only detected in G_0_ at canonical Dicer targets (centromeric, subtelomeric transcripts); y-axis indicates effect size (Δlog_2_(RPM)) between wild-type and *dcr1*Δ quiescent cells (>0 corresponds to Dicer-dependent small RNAs); x-axis indicates strand symmetry (from 0.0 for absolute strand-specificity to 1.0 for equivalent production from both strands; *bona fide* small RNAs typically have a ratio >0.3). (**B**) Overview of the iPARE-seq protocol for specific adapter ligation to available 5’-monophosphate termini, present at cleavage sites and non-RNA pol II transcripts. (**C**) Simplified overview of the main processing sites for maturation of the pre-rRNA into rRNA. The key initial processing steps correspond to cleavage of sites A_0_ and B_0_, balancing co-transcriptional and post-transcriptional processing according to environmental conditions *(26,33)*. iPARE-seq recovers termini with single-nucleotide precision at the mature 18S 5’ end (corresponding to A1 cleavage) as well as pre-rRNA transcription start site, both in cycling cells (**D**) and in quiescent cells (**E**). (**F**) Down-regulation of most 5’ETS cleavage sites and mature rRNA in *dcr1*Δ mutants, and catalytic-dead *dcr1-5* and *rdp1*-D903A. Among these sites, the 28S 5’ is the least affected, and A_0_ site the most affected.

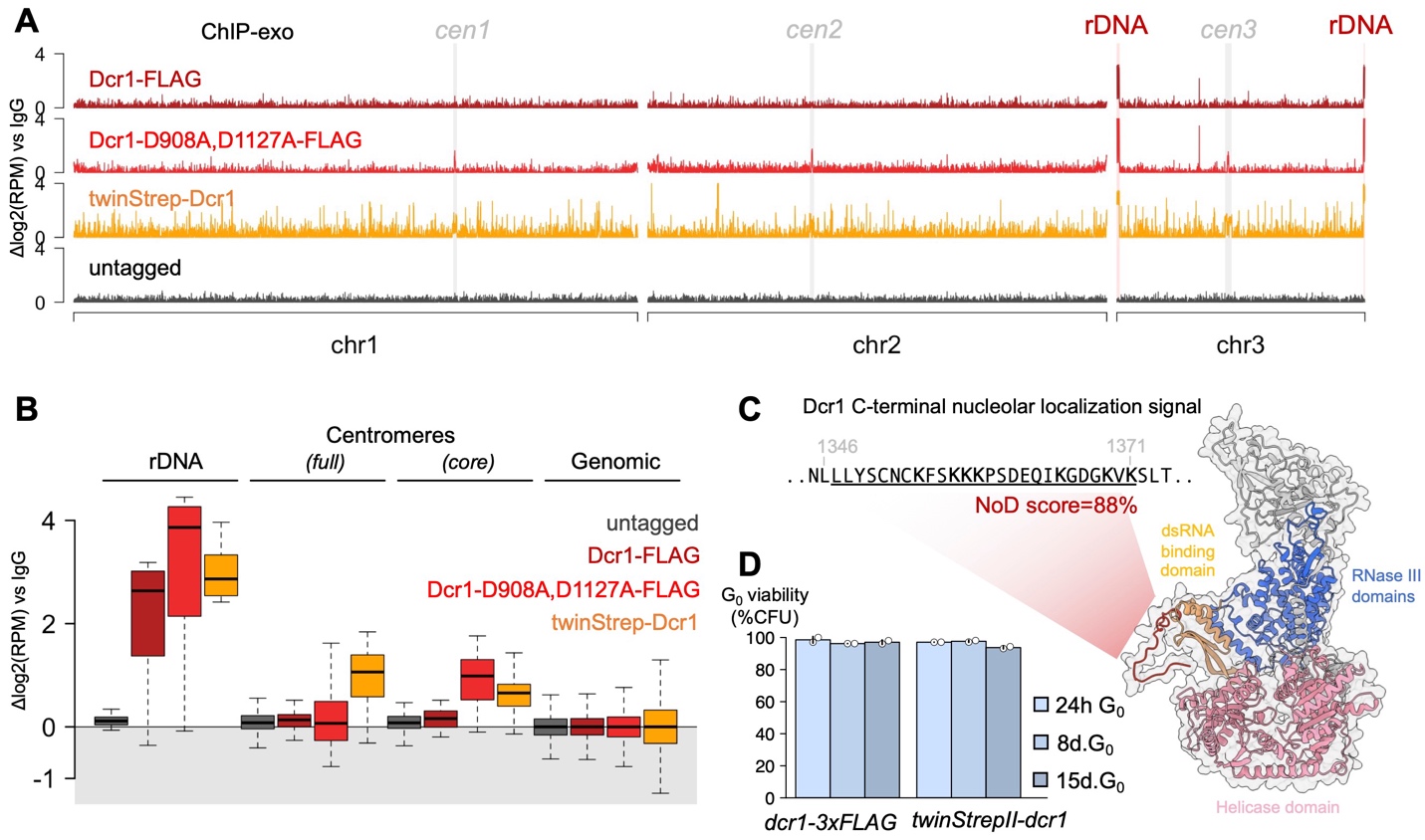

Fig. S2. Dicer binds to ribosomal DNA in quiescent cells.

(**A**) Whole-genome overview of ChIP-exo coverage tracks for C- and N-terminally tagged Dicer constructs. Maximal enrichment is observed specifically at rDNA regions on both ends of chr3. In the catalytic mutant Dcr1-D908A,D1127A-FLAG and the N-terminal tag twinStrepII-Dcr1, enrichment is also seen at centromeric regions. (**B**) Quantification of Dcr1 enrichment at rDNA and centromeres, compared to the rest of the genome. (**C**) Detection of an exposed C-terminal Dicer nucleolar localization signal by NoD *(118)*. (**D**) Tagged Dcr1 strains do not affect its G_0_ function, as seen by wild-type viability over 15 days of quiescence.

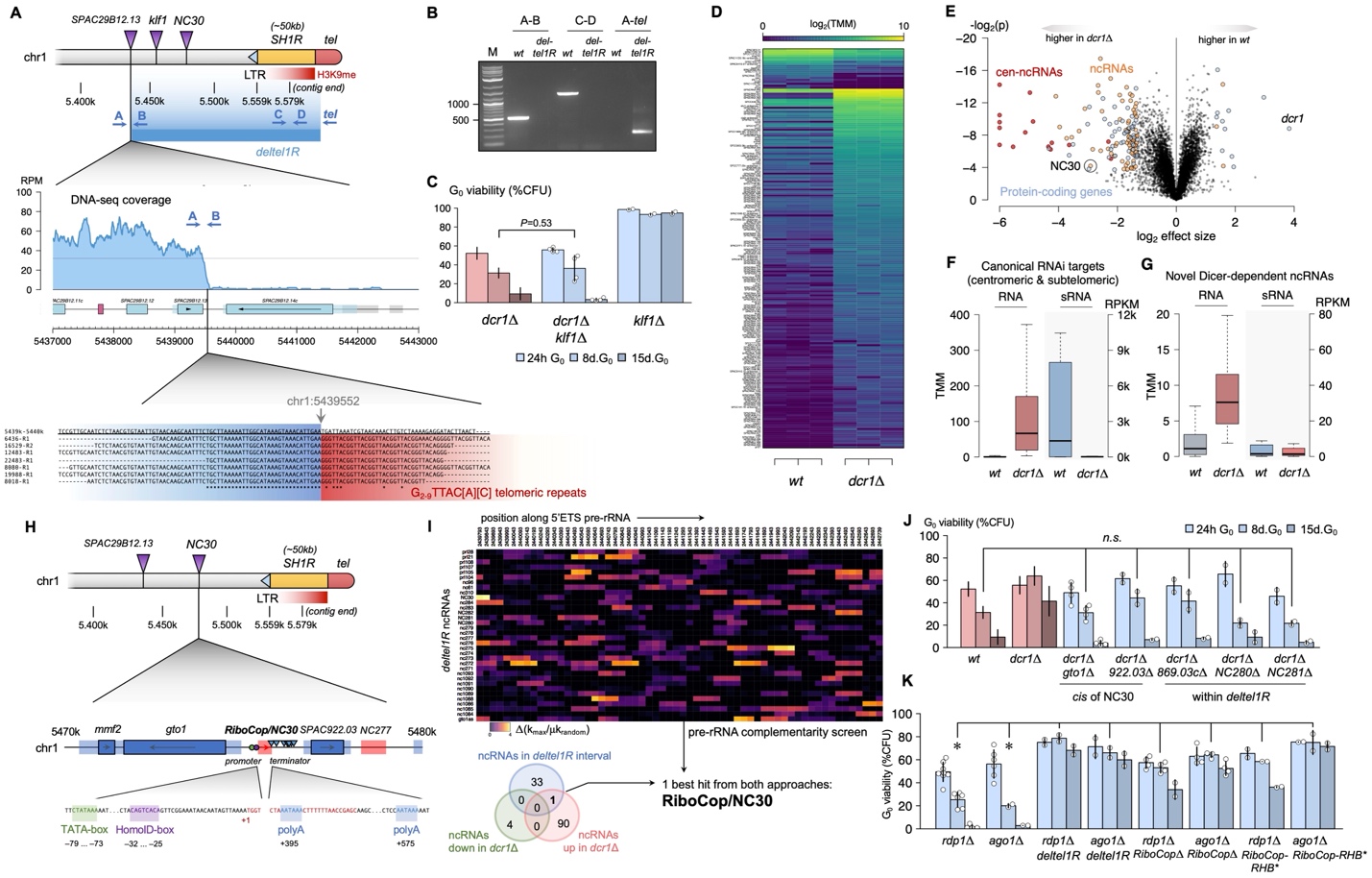

Fig. S3. Identification of *NC30/RiboCop* as the suppressor within the *deltel1R* interval.

(**A**) Location of the *deltel1R* deletion and identification of the breakpoint at chr3:5439552 by DNA-seq. (**B**) PCR confirmation of *deltel1R*. (**C**) Initial candidate gene *klf1* is located in *deltel1R*, but is not responsible for suppression as *dcr1*Δ*klf1*Δ shows the *dcr1*Δ G_0_ phenotype. (**D**) RNA-seq of wt and *dcr1*Δ G0 cells identifies a set of genes, mostly upregulated in *dcr1*Δ; (**E**) most of the upregulated genes correspond to non-coding RNAs, including canonical targets (centromeric RNAs, red dots) and new long non-coding RNAs (lncRNAs, orange dots). (**F**) Canonical RNAi targets are upregulated in *dcr1*Δ and produce Dicer-dependent small RNAs in wild-type. (**G**) Dicer-dependent lncRNAs do not produce significant amount of small RNAs, suggesting that they are indirect Dicer targets. (**H**) Location of the *NC30/RiboCop* ncRNA gene within *deltel1R*, with a ribosomal-type promoter. (**I**) Screen of all ncRNAs within *deltel1R* for potential complementarity regions to the 5’ETS pre-rRNA. The top candidate is the Dicer-regulated ncRNA *NC30/RiboCop*. (**J**) Genes proximal to RiboCop are not causative of the suppression seen in *dcr1*Δ*deltel1R* and *dcr1*Δ*RiboCop*Δ. (**K**) *deltel1R*, *RiboCop*Δ and *RiboCop*-RHB* are not only G_0_ phenotypic suppressors of *dcr1*Δ, but also of *ago1*Δ and *rdp1*Δ.

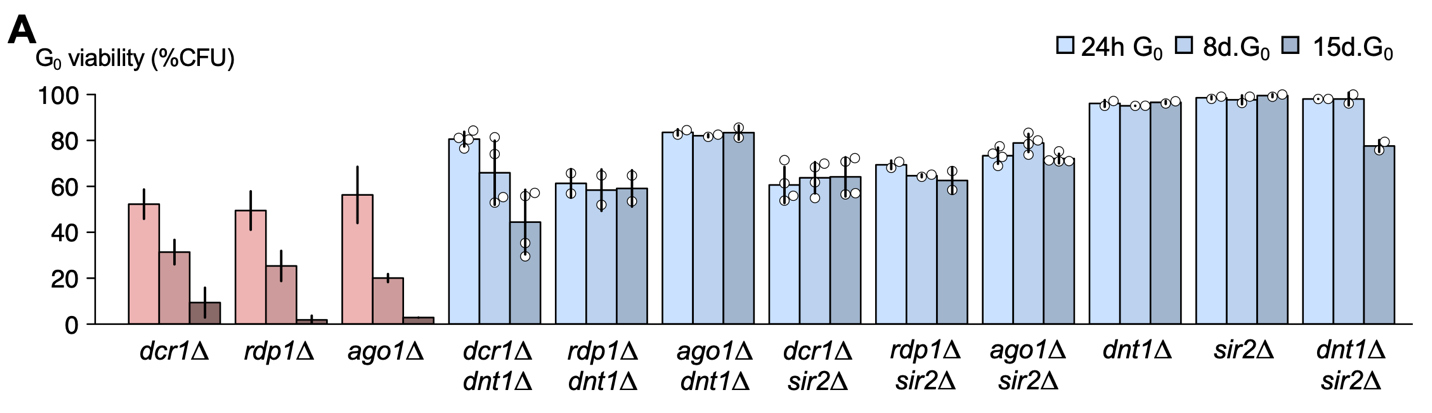

Fig. S4. Genetic suppression of RNAi by RENT mutants in quiescence.

(**A**) Both *dnt1*Δ and *sir2*Δ suppress the loss of viability in G_0_ phenotype of all RNAi mutants (*dcr1*Δ, *rdp1*Δ, *ago1*Δ). The single-mutants *dnt1*Δ, *sir2*Δ, and the *dnt1*Δ*sir2*Δ double-mutant are not affected and survive over 15 days in quiescence.

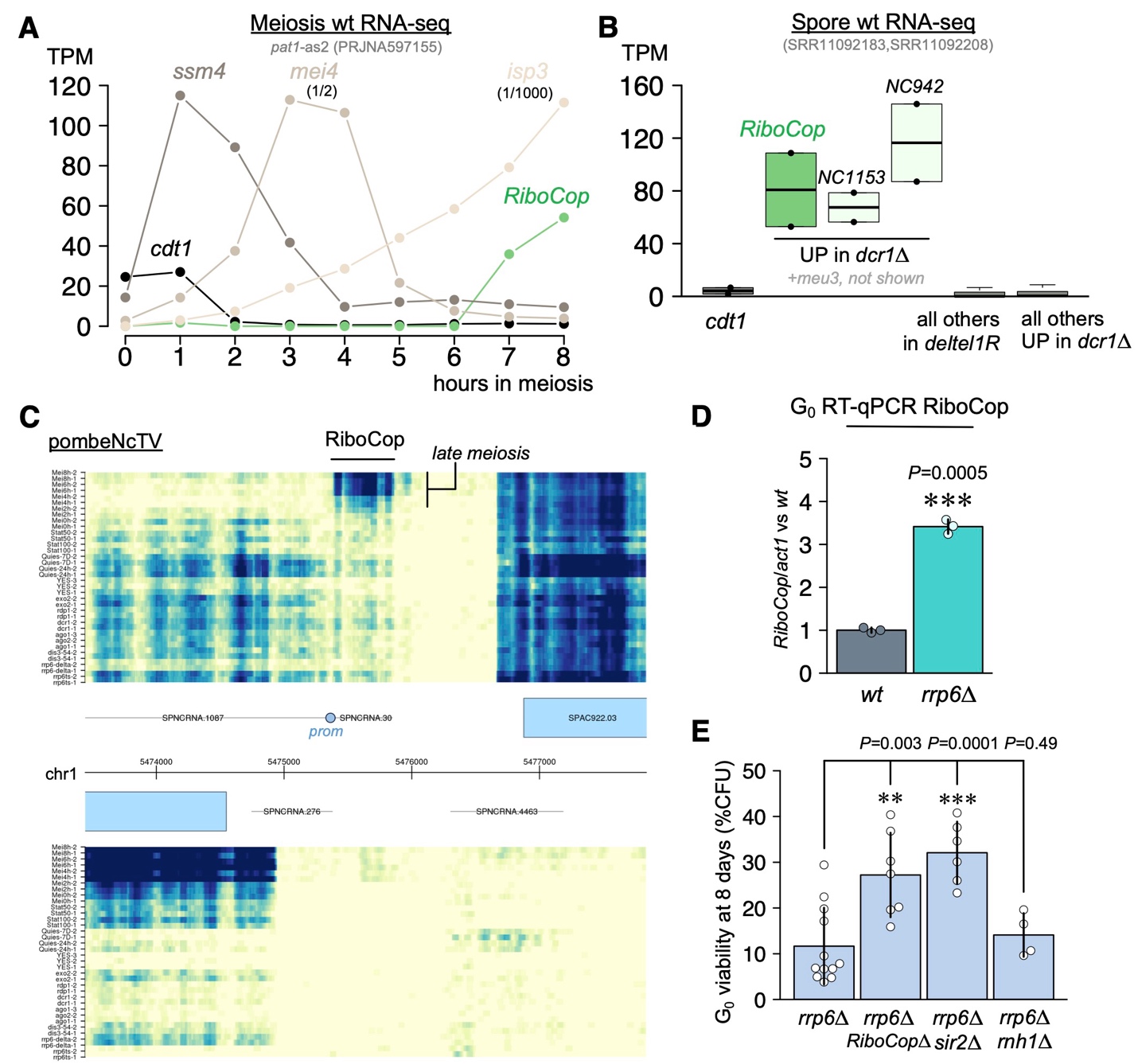

Fig. S5. RiboCop is induced during sporulation and in other pre-rRNA processing mutants.

(**A**) Transcriptome analysis of wild-type *pat1*-as2 mutant *(52)* reveals *RiboCop* expression in late meiosis/sporulation (7-8h). Control genes: *cdt1* (S-phase marker), *ssm4* (early meiosis), *mei4* (middle-phase), *isp3* (late meiosis/sporulation). (**B**) Transcriptome analysis of wild-type spores *(53)* shows high RiboCop expression in dormant spores. RiboCop is the only non-coding RNA within *deltel1R* with this expression pattern, and one of only 4 Dicer-regulated ncRNAs expressed in spores. (**C**) Transcriptome visualization using pombeNcTV *(11)* shows *RiboCop* expression in wild-type late meiosis, and mild de-repression in cycling *dcr1*Δ and *rrp6*-ts mutants. (**D**) RT-qPCR displaying de-repression of RiboCop in *rrp6*Δ mutants at 24h G_0_. (**E**) The low viability in quiescence of *rrp6*Δ (8 days G_0_) is partially suppressed by *RiboCop*Δ and *sir2*Δ, but not *rnh1*Δ, showing that RiboCop is active in *rrp6*Δ and that RiboCop nucleolar stress response is responsible for a significant proportion of quiescence lethality in *rrp6*Δ.

Table S1. Strains used in this study.

List of all strains used in this study, indicating strain number, mating-type, full genotype, corresponding figures when applicable, and origin of allele/mutant. Each strain has a unique identifier and all strains are available upon request to the lead contact.

| **Strain number** | **Genotype** | **Comment** | **Figure** | **Source** |
| --- | --- | --- | --- | --- |
| BR00 | h- |  | Fig 1A | 972 h- |
| BR51 | h+ |  |  | *(15)* |
| BR245 | h- dcr1::dcr1-D908A,D1127A | *dcr1-5* catalytic dead allele | Fig 1A | *(21)* |
| BR371 | h+ dcr1Δ::HphMX |  |  | *(15)* |
| BR522 | h- nuc1::nuc1-(Gly)6-3xFLAG-HphMX |  | Fig 1B | *(15)* |
| BR539 | h- dcr1Δ::KanMX deltel1R | Isolated single colony from suppressor screen | Fig 2B | This study |
| BR547 | h- deltel1R | Back-cross of suppressor (BR51xBR539) | fig S3B | This study |
| BR565 | h- dcr1Δ::KanMX nuc1::nuc1-(Gly)6-3xFLAG-HphMX |  | Fig 1B | *(15)* |
| BR608 | h- dcr1Δ::KanMX |  | Fig 1A | This study |
| BR742 | h- dcr1::dcr1-3xFLAG-HphMX |  | fig S2 | This study |
| BR745 | h- dcr1::dcr1-D908A,D1127A-3xFLAG-HphMX | *dcr1-5* catalytic dead allale | fig S2 | This study |
| BR872 | h+ klf1Δ::HphMX |  |  | This study |
| BR948 | h- klf1Δ::HphMX |  | fig S3C | This study |
| BR950 | h- dcr1Δ::KanMX klf1Δ::HphMX |  | fig S3C | This study |
| BR1009 | h- dcr1Δ::KanMX clr4::NatMX-purg1-clr4 |  | Fig 3E | *(15)* |
| BR1013 | h- clr4::NatMX-purg1-clr4 |  |  | *(15)* |
| BR1046 | h+ dnt1Δ::HphMX |  |  | This study |
| BR1082 | h- rpa2::rpa2-(Gly)6-HphMX |  | Fig 1B | This study |
| BR1102 | h+ dcr1Δ::KanMX dnt1Δ::HphMX |  |  | This study |
| BR1103 | h- dcr1Δ::KanMX dnt1Δ::HphMX |  | fig S4 | This study |
| BR1132 | h- dcr1Δ::KanMX dnt1Δ::HphMX |  | fig S4 | This study |
| BR1170 | h- dcr1Δ::KanMX rpa2::rpa2-(Gly)6-3xFLAG-HphMX |  | Fig 1B | This study |
| BR1197 | h- rpa12::rpa12-3xFLAG-HphMX |  |  | This study |
| BR1211 | h- dcr1Δ::KanMX rpa12::rpa12-3xFLAG-HphMX |  |  | This study |
| BR1270 | h- dnt1Δ::HphMX |  | fig S4 | This study |
| BR1277 | h- dnt1Δ::HphMX nuc1::nuc1-(Gly)6-3xFLAG-HphMX |  |  | This study |
| BR1345 | h+ dcr1Δ::KanMX |  |  | This study |
| BR1349 | h- NC30Δ::NatMX |  | Fig 2G, Fig 3 | This study |
| BR1352 | h- NC280Δ::NatMX |  |  | This study |
| BR1355 | h- NC281Δ::NatMX |  |  | This study |
| BR1364 | h- dcr1Δ::KanMX NC30Δ::NatMX |  | Fig 2B | This study |
| BR1366 | h- dcr1Δ::KanMX NC280Δ::NatMX |  | fig S3J | This study |
| BR1368 | h- dcr1Δ::KanMX NC281Δ::NatMX |  | fig S3J | This study |
| BR1383 | h- dcr1Δ::KanMX NC30Δ::NatMX | BR1349 transformed with dcr1Δ |  | This study |
| BR1384 | h- dcr1Δ::KanMX NC30Δ::NatMX | BR1349 transformed with dcr1Δ |  | This study |
| BR1387 | h- dcr1Δ::KanMX NC30Δ::NatMX | Independent cross (BR1345xBR1349) |  | This study |
| BR1388 | h- dcr1Δ::KanMX NC30Δ::NatMX | Independent cross |  | This study |
| BR1415 | h- gto1Δ::NatMX |  |  | This study |
| BR1417 | h- SPAC922.03Δ::NatMX |  |  | This study |
| BR1419 | h- SPAC869.03cΔ::NatMX |  |  | This study |
| BR1421 | h- yhb1Δ::NatMX |  |  | This study |
| BR1441 | h- dcr1Δ::KanMX gto1Δ::NatMX |  | fig S3J | This study |
| BR1445 | h- dcr1Δ::KanMX SPAC922.03Δ::NatMX |  | fig S3J | This study |
| BR1447 | h- dcr1Δ::KanMX SPAC869.03cΔ::NatMX |  | fig S3J | This study |
| BR1450 | h- dcr1Δ::KanMX yhb1Δ::NatMX |  | fig S3J | This study |
| BR1472 | h- NC30Δ::NatMX nuc1::nuc1-(Gly)6-3xFLAG-HphMX |  |  | This study |
| BR1475 | h- dcr1Δ::KanMX NC30Δ::NatMX nuc1::nuc1-(Gly)6-3xFLAG-HphMX |  |  | This study |
| BR1478 | h- dcr1Δ::HphMX NC30Δ::NatMX | Independent dcr1Δ mutant (BR371xBR1349) | Fig 2B | This study |
| BR1480 | h- NC30Δ::NatMX otr1R(SphI)::ade6+ imr1L(NcoI)::ura4+ ura4-DS/R ade6-DN/N leu1-32 | | Fig 3C | This study |
| BR1487 | h- dcr1Δ::KanMX deltel1R nuc1::nuc1-(Gly)6-3xFLAG-HphMX |  |  | This study |
| BR1490 | h- deltel1R nuc1::nuc1-(Gly)6-3xFLAG-HphMX |  |  | This study |
| BR1492 | h+ (gto1,NC30,SPAC922.03)Δ::NatMX |  |  | This study |
| BR1495 | h- (gto1,NC30,SPAC922.03)Δ::NatMX |  |  | This study |
| BR1500 | h- dcr1Δ::KanMX (gto1,NC30,SPAC922.03)Δ::NatMX |  |  | This study |
| BR1606 | h- rdp1Δ::KanMX deltel1R |  | fig S3K | This study |
| BR1623 | h- deltel1R IGR(arg3--arg11)::NC30(+IGRs)-HphMX |  |  | This study |
| BR1638 | h- dcr1Δ::KanMX deltel1R IGR(arg3--arg11)::NC30(+IGRs)-HphMX |  | Fig 2C | This study |
| BR1639 | h- dcr1Δ::KanMX deltel1R IGR(arg3--arg11)::NC30(+IGRs)-HphMX |  | Fig 2C | This study |
| BR1646 | h- dcr1Δ::KanMX IGR(arg3--arg11)::NC30(+IGRs)-HphMX |  | Fig 2C | This study |
| BR1657 | h- deltel1R IGR(arg3--arg11)::NC30-RHB*(+IGRs)-HphMX |  |  | This study |
| BR1665 | h+ dcr1Δ::KanMX deltel1R |  |  | This study |
| BR1670 | h- arg3Δ::KanMX |  |  | This study |
| BR1677 | h- ago1Δ::KanMX NC30Δ::NatMX |  | fig S3K | This study |
| BR1681 | h- rdp1Δ::KanMX NC30Δ::NatMX |  | fig S3K | This study |
| BR1684 | h- rdp1Δ::KanMX NC30::HphMX-NC30-RHB* |  | fig S3K | This study |
| BR1685 | h- ago1Δ::KanMX deltel1R |  | fig S3K | This study |
| BR1694 | h- ago1Δ::KanMX NC30::HphMX-NC30-RHB* |  | fig S3K | This study |
| BR1701 | h- dcr1Δ::KanMX deltel1R IGR(arg3--arg11)::NC30-RHB*(+IGRs)-HphMX | | Fig 2C | This study |
| BR1713 | h+ NC30::HphMX-NC30 |  |  | This study |
| BR1733 | h- NC30::HphMX-NC30 |  |  | This study |
| BR1736 | h- dcr1Δ::KanMX NC30::HphMX-NC30 |  |  | This study |
| BR1738 | h+ NC30::HphMX-NC30-RHB* |  |  | This study |
| BR1766 | h+ rrp6Δ::HphMX |  |  | This study |
| BR1773 | h- dcr1Δ::KanMX NC30::HphMX-NC30-RHB* |  | Fig 2B | This study |
| BR1787 | h- dcr1Δ::KanMX rpa12Δ::HphMX deltel1R IGR(arg3--arg11)::NC30(+IGRs)-HphMX | | Fig 3D | This study |
| BR1792 | h- dcr1Δ::KanMX swi6-W293* deltel1R IGR(arg3--arg11)::NC30(+IGRs)-HphMX | | Fig 3D | This study |
| BR1804 | h- deltel1R IGR(leu2)::NC30(+IGRs)-HphMX |  |  | This study |
| BR1810 | h- dcr1Δ::KanMX deltel1R IGR(leu2)::NC30(+IGRs)-HphMX |  | Fig 2C | This study |
| BR1854 | h- dcr1Δ::KanMX dnt1Δ::HphMX nuc1::nuc1-(Gly)6-3xFLAG-HphMX |  |  | This study |
| BR1889 | h- rdp1::rdp1-D903A-KanMX |  | fig S1F | This study |
| BR2000 | h+ sir2Δ::NatMX |  |  | This study |
| BR2016 | h- sir2Δ::NatMX |  | fig S4 | This study |
| BR2018 | h- dcr1Δ::KanMX sir2Δ::NatMX |  | fig S4 | This study |
| BR2040 | h- dnt1Δ::HphMX sir2Δ::NatMX |  | fig S4 | This study |
| BR2091 | h- dcr1::KanMX-twinStrepII-dcr1 |  | fig S2 | This study |
| BR2114 | h- dcr1Δ::KanMX dnt1Δ::HphMX deltel1R IGR(arg3--arg11)::NC30(+IGRs)-HphMX | | Fig 3D | This study |
| BR2116 | h- dcr1Δ::KanMX rpa12Δ::HphMX deltel1R IGR(arg3--arg11)::NC30(+IGRs)-HphMX | | Fig 3D | This study |
| BR2118 | h- dcr1Δ::KanMX clr4Δ::KanMX deltel1R IGR(arg3--arg11)::NC30(+IGRs)-HphMX | | Fig 3D | This study |
| BR2121 | h- dcr1Δ::KanMX sir2Δ::NatMX deltel1R IGR(arg3--arg11)::NC30(+IGRs)-HphMX | | Fig 3D | This study |
| BR2148 | h- NC30Δ::NatMX clr4::NatMX-purg1-clr4 |  |  | This study |
| BR2149 | h- dcr1Δ::KanMX NC30Δ::NatMX clr4::NatMX-purg1-clr4 |  | Fig 3E | This study |
| BR2150 | h- dcr1Δ::KanMX sir2Δ::NatMX clr4::NatMX-purg1-clr4 |  | Fig 3E | This study |
| BR2240 | h- sir2Δ::NatMX clr4::NatMX-purg1-clr4 |  |  | This study |
| BR2243 | h- dnt1Δ::HphMX clr4::NatMX-purg1-clr4 |  |  | This study |
| BR2249 | h- rrp6Δ::HphMX |  | fig S5DE | This study |
| BR2298 | h- NC30::HphMX-NC30-RHB* |  |  | This study |
| BR2301 | h- rdp1Δ::KanMX NC30::HphMX-NC30-RHB* |  | fig S3K | This study |
| BR2304 | h- ago1Δ::KanMX NC30::HphMX-NC30-RHB* |  | fig S3K | This study |
| BR2307 | h- rdp1Δ::KanMX dnt1Δ::HphMX |  | fig S4 | This study |
| BR2310 | h- ago1Δ::KanMX dnt1Δ::HphMX |  | fig S4 | This study |
| BR2317 | h- rdp1Δ::KanMX sir2Δ::NatMX |  | fig S4 | This study |
| BR2322 | h- rrp6Δ::HphMX NC30Δ::NatMX |  | fig S5E | This study |
| BR2367 | h- ago1Δ::KanMX sir2Δ::NatMX |  | fig S4 | This study |
| BR2368 | h+ pabp::pabp-(Gly)6-3xFLAG-HphMX |  |  | This study |
| BR2369 | h+ enp2::enp2-(Gly)6-3xFLAG-HphMX |  |  | This study |
| BR2371 | h+ rnh1::rnh1-(Gly)6-3xFLAG-HphMX |  |  | This study |
| BR2385 | h- dcr1Δ::KanMX enp2::enp2-(Gly)6-3xFLAG-HphMX |  | Fig 4D | This study |
| BR2388 | h- dcr1Δ::KanMX rnh1::rnh1-(Gly)6-3xFLAG-HphMX |  | Fig 4D | This study |
| BR2390 | h- dcr1Δ::KanMX NC30Δ::NatMX rnh1::rnh1-(Gly)6-3xFLAG-HphMX |  | Fig 4D | This study |
| BR2391 | h- rnh1Δ::HphMX |  |  | This study |
| BR2393 | h- rnh1::rnh1-HBDΔ-HphMX | NTD/HBD domain deletion |  | This study |
| BR2398 | h- dcr1Δ::KanMX pabp::pabp-(Gly)6-3xFLAG-HphMX | SPAC57A7.04c |  | This study |
| BR2403 | h- rnh1::rnh1-D191N-HphMX | Rnh1 catalytic mutant |  | This study |
| BR2433 | h- dcr1Δ::KanMX rnh1Δ::HphMX |  | Fig 4F | This study |
| BR2444 | h- dcr1Δ::KanMX rnh1::rnh1-HBDΔ-HphMX |  | Fig 4F | This study |
| BR2447 | h- dcr1Δ::KanMX rnh1::rnh1-D191N-HphMX |  | Fig 4F | This study |
| BR2495 | h+ enp2::enp2-E521Δ-HphMX ade6-M216 | Sporulated from ade6-M210/ade6-M216 diploid | | This study |
| BR2497 | h+ enp2::enp2-P486Δ-HphMX ade6-M216 | Sporulated from ade6-M210/ade6-M216 diploid | | This study |
| BR2499 | h+ enp2::enp2-L465Δ-HphMX ade6-M210 | Sporulated from ade6-M210/ade6-M216 diploid | | This study |
| BR2500 | h+ rnt1::purg1-rnt1-NatMX |  |  | This study |
| BR2503 | h- enp2::enp2-E521Δ-HphMX |  |  | This study |
| BR2507 | h- enp2::enp2-P486Δ-HphMX |  | Fig 4F | This study |
| BR2510 | h- rnt1::purg1-rnt1-NatMX |  |  | This study |
| BR2511 | h+ dcr1Δ::KanMX rnt1::purg1-rnt1-NatMX |  |  | This study |
| BR2512 | h- dcr1Δ::KanMX rnt1::purg1-rnt1-NatMX |  | Fig 1A | This study |
| BR2515 | h- dcr1Δ::KanMX enp2::enp2-E521Δ-HphMX |  |  | This study |
| BR2518 | h- dcr1Δ::KanMX enp2::enp2-P486Δ-HphMX |  |  | This study |
| BR2520 | h- rrp6Δ::HphMX sir2Δ::NatMX |  | fig S5E | This study |
| BR2522 | h- enp2::enp2-L465Δ-HphMX |  | Fig 4F | This study |
| BR2525 | h- rrp6Δ::HphMX rnh1Δ::NatMX |  | fig S5E | This study |
| HSK351 | h+ Δclr4:KanMX otr1R(SphI)::ade6+ imr1L(NcoI)::ura4+ ura4-DS/R ade6-DN/N leu1-32 | | Fig 3C | *(119)* |
| FY2002 | h+ otr1R(SphI)::ade6+ imr1L(NcoI)::ura4+ ura4-DS/R ade6-DN/N leu1-32 | | Fig 3C | *(120)* |

Table S2. Primers used in this study.

List of all primers used in this study, with sequence and usage.

| **Primer name** | **Primer sequence** | **Application** |
| --- | --- | --- |
| bPAR3OH | /5Biosg/rUrCrUrArGrArArUrGrCrArUrGrGrGrCrCrCrUrCrCrArArG | iPARE-seq 5' RNA adapter |
| PARss | NNNNTCTAGAATGCATGGGCCCTCCAAG | iPARE-seq second-strand synthesis primer |
| N1-rnt1 | TTCATATTGACGGTTTGGTAGTT | p.urg1-rnt1 (G0 overexpression) *(32)* |
| N2-rnt1 | TAATTAACCCGGGGATCCGTCGACCTCAATGTTATAATCTTCAAATCTACCAA |  |
| L5-purg1-800 | AAACGAGCTCGAATTCATCGATGATATTAGTTCAGCTTGTATACATTGATTCC |  |
| purg1-rev | ATTGAATTAGTTCTAATTTAGTAATTGAAGAAAAC |  |
| V2-purg1-rnt1 | GTTTTCTTCAATTACTAAATTAGAACTAATTCAATATGGGACGGTTTAAGAGGCATCAT |  |
| N6-rnt1 | ATTTCGTAGACTCCTTCAATATGG |  |
| rnt1-qPCR-forw | CAAATCAGTCGAATCCGAATGA | Rnt1 qPCR |
| rnt1-qPCR-rev | GAAAGACTTCCTTCATCCATTTGA |  |
| dcr1-qPCR-forw | GGCTAAAGGAGATATTGAACACAAG | Dcr1 qPCR |
| dcr1-qPCR-rev | GCAACTTTACGGGATTTGCC |  |
| actin-qPCR-forw | TGCACCTGCCTTTTATGTTG | act1 qPCR |
| actin-qPCR-rev | TGGGAACAGTGTGGGTAACA |  |
| 18S-13-forw | CCTGCCAGTAGTCATATGCTTG | 18S rDNA qPCR |
| 18S-253-rev | GACTCACCAAAAAAAGCCCG |  |
| p30-qPCR-forw | CCATATCAATTTCCCATGTTCC | centromere qPCR |
| p30-qPCR-rev | CATCAAGCGAGTCGAGATGA |  |
| L3-rdp1 | GTGTAGCTGATGAAACTGGAACA | Rdp1 catalytic mutant D903A (GAT->GCA) |
| rdp1-D903A-MF | GTGATCTGGATGGGGcaGAGTACA |  |
| rdp1-D903A-MR | TATAACTGTGTACTCtgcCCCATCCAG |  |
| L10-rdp1 | TAATTAACCCGGGGATCCGTCGACCAAATTGCTTTAACCATCGAGTATATGT |  |
| L11-rdp1 | AAACGAGCTCGAATTCATCGATGATAAATCGTAATAATGTATAATTAACGATTTTTATATC |  |
| L12-rdp1 | CTCATTCGAAAGCTAGCGAGT |  |
| N1-dcr1 | GAATACTTTTCATTTTAGAGACTTCAAATAATA | Tagging of Dcr1 |
| N2-dcr1 | TAATTAACCCGGGGATCCGTCGACCCCTGCTCTTATCGGAACATAGGAAAA |  |
| N5-dcr1 | AAACGAGCTCGAATTCATCGATGATACAGTGAACGCGTAAACAGATATATATG |  |
| NF-twinStrep-dcr1 | CTTTATTTTCTTTTTGAATGgccagcgcttggagccacccgcagttcgagaaaggtggaggttccggaggtggatcgggaggttcggcgtggagccacccgcagttcgaaaaaggcggaggaGATATTTCAAGTTTTCTACTTCCTCAACT | |
| NR-twinStrep-dcr1 | CGGGTGGCTCCAAGCGCTGGCCATTCAAAAAGAAAATAAAGGCG |  |
| N6-dcr1 | GATCGAATGTATTCCGCTTGTT |  |
| L3C-rpa2 | GTGGGTCTATTATTTCTATCATGTCTAC | Tagging rpa2 |
| L4C-Gly6-rpa2 | TAATTAACCCGGGGATCCGTCGACCTCCACCTCCTCCACCACCTTTAACTTCAAGCATCATTTTAATATTC | |
| L5-rpa2 | AAACGAGCTCGAATTCATCGATGATAATTAAAGTGTTGAAAAGAGTATTGAAATTA |  |
| L6-rpa2 | CCACCTAGACCTACCTTTATAGTTTTC |  |
| del-A | GACGCACAAGTTTGTATAGACGAC | Confirmation of deltel1R |
| del-B | GGAAAGTCAGGAGGTATCGATATC |  |
| del-C | GGTACATGTACATCGTTGTCTTATGC |  |
| del-D | GAAATCTACGACCATTTGACCC |  |
| telom-rev | AACCGTAACCCCTGTAACCGTA |  |
| SPAC750.02c-qPCR-forw | GCCAAGTATTTCCAATATTGCAG | Confirmation of deltel1R by DNA-qPCR (CNV) |
| SPAC750.02c-qPCR-rev | CGCTATAACCAATTAAATATATAGGTCTT |  |
| ftm4-qPCR-forw | CGAGACCAATGACAATGGAT |  |
| ftm4-qPCR-rev | TAGTACTCCTCCCATTTCAGCTAC |  |
| L1-NC30 | CGTAACATTAGAACGGTTCTATCCA | Deletion of NC30 |
| L2-NC30 | TAATTAACCCGGGGATCCGTCGACCTTTTAACTATTGTTATTTCCGAACTGT |  |
| L5-NC30 | AAACGAGCTCGAATTCATCGATGATAAAGCTTTGTTAGTGTATACAAATCGAT |  |
| L6-NC30 | ATTCTCAATGGTATGTAAGGTATTACTAAT |  |
| NC30-chk-forw | CCTAGGACGGATATACATTACTCTC |  |
| NC30-chk-rev | CAGTTTAGCAAGTTTACATGTATTCTG |  |
| NC30-qPCR-forw | GATGGTGACATTTTTTGCGATAAC | NC30 RT-qPCR (with chk-rev) |
| L1-NC280 | CGGAATAGAAGATTGAAATAACTACA | Deletion of NC280 |
| L2-NC280 | TAATTAACCCGGGGATCCGTCGACCGTCTTGTTGTATCAACCTTCAAGGT |  |
| L5-NC280 | AAACGAGCTCGAATTCATCGATGATAATCTTTAATATTTGCATTTGAACTTACA |  |
| L6-NC280 | GTGGATCATCGCTTACTTAAAACT |  |
| NC280-chk-forw | CTAAGGAAGTAGCGGTTGTCAAA |  |
| L1-NC281 | CTAAGCTGAGAAGTACGTTTTATTGAAG | Deletion of NC281 |
| L2-NC281 | TAATTAACCCGGGGATCCGTCGACCTATTGCAAGGGTTTACAATTATTCCT |  |
| L5-NC281 | AAACGAGCTCGAATTCATCGATGATAGATACCAATGCATCACTATGTTAAATA |  |
| L6-NC281 | TGCCAAAACAATTATCTGTTTAATG |  |
| NC281-chk-forw | CTTACATTGACTAGGTTCTTCAACA |  |
| NC281-chk-rev | CCCTCTTACTTATGCACAGATCA |  |
| L1-gto1 | GAGCTTTGAGATATCTAGCATTCC | Deletion of gto1 |
| L2-gto1 | TAATTAACCCGGGGATCCGTCGACCAGTTATCAATAAAAAAAACTAGAAGGTGG |  |
| L5-gto1 | AAACGAGCTCGAATTCATCGATGATAATGACACAATTTGTTTAATTACTGATTTT |  |
| L6-gto1 | GGTGGTAAAATTGAAATCGAATG |  |
| gto1-chk-forw | GGATCATGTGGAACAGATAAACTAGA |  |
| gto1-chk-rev | GATTGTATCTAACTGTTCCGTCATGG |  |
| L1-SPAC922.03 | GGCGTTTAATACAACCTCAGGAA | Deletion of SPAC922.03 |
| L2-SPAC922.03 | TAATTAACCCGGGGATCCGTCGACCTGTAATGTTTGTTAAACGAGACTGAATAA |  |
| L5-SPAC922.03 | AAACGAGCTCGAATTCATCGATGATATTACCGCTAACTATTAATACTTTTAATATTTAGAG |  |
| L6-SPAC922.03 | CAACGATGCTATAGATGCAATC |  |
| SPAC922.03-chk-forw | TCTTGAAGAGCTTACAAAGAAGG |  |
| SPAC922.03-chk-rev | GACATCGCTCTCAGTCAATTC |  |
| L1-SPAC869.03c | GCCTCAATATGTTGTACCTTATGC | Deletion of SPAC869.03c |
| L2-SPAC869.03c | TAATTAACCCGGGGATCCGTCGACCTTTGAGTTTCCAAATAATAGTCTAAGTTCC |  |
| L5-SPAC869.03c | AAACGAGCTCGAATTCATCGATGATAATCTTAACTTAAATGACAAGCTTTGCAG |  |
| L6-SPAC869.03c | AATGAGGACTAAATTTGCGCAA |  |
| SPAC869.03c-forw | GTATGCCTTTACTATATGGAATGACTG |  |
| SPAC869.03c-rev | CAACACCTGTGATAGATCCCAT |  |
| L1-yhb1 | CTACCTGAATGGCTTCTGAGT | Deletion of yhb1 |
| L2-yhb1 | TAATTAACCCGGGGATCCGTCGACCTTTACTTTGGAATCTCTAATTCCAA |  |
| L5-yhb1 | AAACGAGCTCGAATTCATCGATGATAGTCGCAAATGCAGCAAAAT |  |
| L6-yhb1 | TCATTATCCATAGTCACAAATAGCAA |  |
| yhb1-forw | AGAGGCATGGACTACTGCAT |  |
| yhb1-rev | TGACACCAACGATATCACCAA |  |
| L1-klf1 | CCCTGTATAATAGCCAGAAGGTT | Deletion of klf1 |
| L2-klf1 | TAATTAACCCGGGGATCCGTCGACCGAAGAAACGCAAGTAACCTCAC |  |
| L5-klf1 | AAACGAGCTCGAATTCATCGATGATACTTGAGGATGTATAACGAACGATT |  |
| L6-klf1 | GGATTTATTTTGAAATTAAGGACCAT |  |
| klf1-forw | GTTCGCATTTTAGAGTTGACGA |  |
| klf1-rev | CACCATGAAAAATTTGAGAGAGG |  |
| arg3-11-IGR-L1 | CTAGTCCTAACTGACACAGTACAATATTCAT | Amplification of IGR(arg3--arg11) region for transgene insertion |
| arg3-11-IGR-NC30-R | AACCACCTTCTAGTTTTTTTTATTGATAACTAACGTCAGAATACTTATCAGCAATAGCTT |  |
| arg3-11-IGR-L5 | AAACGAGCTCGAATTCATCGATGATACTGTATCAATATTACCATGAATAACATAAG |  |
| arg3-11-IGR-L6 | GATATTCAGGGTGAGATTGCATA |  |
| NC30-IGR-fw | AGTTATCAATAAAAAAAACTAGAAGGTGGTT | Amplification of NC30 including full IGRs on either side |
| NC30-IGR-rev | TAATTAACCCGGGGATCCGTCGACCTGTAATGTTTGTTAAACGAGACTGAA |  |
| leu2-IGR-L1 | TTACATATTGTCTGCTTAATAATCGATATAA | Amplification of IGR(leu2) region for transgene insertion |
| leu2-IGR-NC30-R | AACCACCTTCTAGTTTTTTTTATTGATAACTCCTTATCAGCTGTATTATCGAGTTATAC |  |
| leu2-IGR-L5 | AAACGAGCTCGAATTCATCGATGATACTATTATAGTGTTTCATACCCTGCATTA |  |
| leu2-IGR-L6 | GTAGATGGTATTGATACCACGGAA |  |
| L10-NC30 | TAATTAACCCGGGGATCCGTCGACCTATATTTATTCTCAATGGTATGTAAGGTATTAC | NC30 mutation (marker downstream) |
| L11-NC30 | AAACGAGCTCGAATTCATCGATGATAGAAACAAAAATAAATAAATAAACAAGATTACTAAA | |
| L12-NC30 | CAATACTTACAACGTACAAGTTAGCAC |  |
| NC30-RHB*-MF | AGTTTCaaacttggaaCCTAGGACGGATATAC | Mutation of NC30 RHB box |
| NC30-RHB*-MR | GttccaagtttGAAACTCCTCGGCTTT |  |
| L1-rrp6 | CTATAGTTGCCTGCTAATAGTTTCACA | Deletion of rrp6 |
| L2-rrp6 | TAATTAACCCGGGGATCCGTCGACCTATGCTATTGTTTACCCTACTGGATCA |  |
| L5-rrp6 | AAACGAGCTCGAATTCATCGATGATATATCAGTTTTAATTTTAAATAAACGAAACAA |  |
| L6-rrp6 | GACACGAATCTTTACGACATGCT |  |
| rrp6-forw | GCTCAATCTGATACGCATTATTTGT |  |
| rrp6-rev | CTTAGAAATTGAAAACACATCAGCC |  |
| L1-dnt1 | GTTCATCACTGCTGTCCCA | Deletion of dnt1 |
| L2-dnt1 | TAATTAACCCGGGGATCCGTCGACCTTCGAATATTCAGCAAATTTCTTTA |  |
| L5-dnt1 | AAACGAGCTCGAATTCATCGATGATAATTTCTTGTTTTTGATTAACTCCCT |  |
| L6-dnt1 | GAACGTTCCATTTAGAATTTCAAC |  |
| dnt1-forw | CTCTCCTGGATTTGTATACAGACC |  |
| dnt1-rev | TCAATTCCATCAATGGTTTCA |  |
| L1-sir2 | GCTTCTAATAGCCATTATTACCAGGAAAT | Deletion of sir2 |
| L2-sir2 | TAATTAACCCGGGGATCCGTCGACCGACAGAGTTTGTATAACATGAAAATATGACTA |  |
| L5-sir2 | AAACGAGCTCGAATTCATCGATGATATTTTAGGACTGATCGATCTATCGAAGTT |  |
| L6-sir2 | ACTAATTACACAGTCCTTGGTATTGTAAAAT |  |
| sir2-forw | CTGCTATCGATATACTAACGTGCTAATTAG |  |
| sir2-rev | GGCAAAAGATCTCTAGCAAAAGTATAA |  |
| L1-arg3 | CTAGCTTGTTTGCAACTGAACTCTA | Deletion of arg3 |
| L2-arg3 | TAATTAACCCGGGGATCCGTCGACCTTTCAATTGCAAACTGAATAATTCT |  |
| L5-arg3 | AAACGAGCTCGAATTCATCGATGATAACTGAAATCCTTAATTTCATTTTTATCTG |  |
| L6-arg3 | CACAAAGCTTGTTGGACTAATGAG |  |
| arg3-forw | GTTTCTGGTATTGTTGCTCGC |  |
| arg3-rev | CAACCTTGGGGTCGTTAACAAT |  |
| L1-rnh1 | TGTATCGTACTGCAGTAGTCCAGATATG | Deletion of rnh1 |
| L2-rnh1 | TAATTAACCCGGGGATCCGTCGACCTTCGTAATGCGAAGATTCAAAAATAG |  |
| L5-rnh1 | AAACGAGCTCGAATTCATCGATGATAATGGTCATCTATGATGAGCGCT |  |
| L6-rnh1 | GCGAAGTGTCATTTTCAACGTT |  |
| N1-rnh1 | AGCACAGCTCTAAATGATATATACAGC | Proximal marker for Rnh1 mutations |
| N2-rnh1 | TAATTAACCCGGGGATCCGTCGACCTTGGCCTCTGAGTAGTTTGCAAAT |  |
| N3-rnh1 | AAACGAGCTCGAATTCATCGATGATACAGAGATGAGGATTCTTTATGAAAGT |  |
| rnh1-D191N-MF | CCATCCGCTCTaagAGCAATTATTCCATTAAGT | Rnh1 catalytic mutant |
| rnh1-D191N-MR | GAATAATTGCTcttAGAGCGGATGGTCAAATC |  |
| rnh1-N-DF | ATGGGTGGAAATAAGCGTGCACAGGAATTTTGCAGGACCGAA | Rnh1 NTD/HBD domain deletion |
| rnh1-N-DR | GGTCCTGCAAAATTCCTGTGCACGCTTATTTCCACCCATTTC |  |
| L1-enp2 | GGCAATGATGAGACCTCAGTTCTTT | Deletion of enp2 |
| L2-enp2 | TAATTAACCCGGGGATCCGTCGACCTCCGTTGAATTCAACTCGGAAAAAATCA |  |
| L5-enp2 | AAACGAGCTCGAATTCATCGATGATA |  |
| L6-enp2 | TGCCAGATTGCAACAAGAAGTAAT |  |
| L3C-enp2 | CTGATGAGGAGTCACTGTCTGATATG | Tagging of enp2 |
| L4C-Gly6-enp2 | TAATTAACCCGGGGATCCGTCGACCTCCACCTCCTCCACCACC/CATGTTTCGGAAAACGTTTTTG | |
| L7-enp2 | AAACGAGCTCGAATTCATCGATGATATTATGATTACTTCTTGTTGCAATCTGG | Truncations of enp2 |
| L8-enp2 | CTTTTAAAACTTGCTTTACTGTCTTAGAGA |  |
| enp2-D-forw | ACAGCTTTAAAATATAGGAACGATGGACT |  |
| enp2-E521-MF | TATAAGCAATTACATCCTTCTAGGTCTtagCCAAAGCACGGTAGAACTGTCAAATT |  |
| enp2-E521-MR | AATTTGACAGTTCTACCGTGCTTTGGctaAGACCTAGAAGGATGTAATTGCTTATA |  |
| enp2-P486-MF | GAAAACTGGCGTTACGGACGGAtagCCAAAGCACGGTAGAACTGTCAAATT |  |
| enp2-P486-MR | AATTTGACAGTTCTACCGTGCTTTGGctaTCCGTCCGTAACGCCAGTTTTC |  |
| enp2-L465-MF | GAATCGTCCAAAAGTCAATGCTGGAtaaCCAAAGCACGGTAGAACTGTCAAATT |  |
| enp2-L465-MR | AATTTGACAGTTCTACCGTGCTTTGGttaTCCAGCATTGACTTTTGGACGATTC |  |
| enp2-D-rev | TAATTAACCCGGGGATCCGTCGACCCATAAAACAATGATAGATACATAAAGTAGATA |  |
| L3C-pabp | CGTCCTATGATGATGCCTGGT | Tagging of PABP |
| L4C-Gly6-pabp | TAATTAACCCGGGGATCCGTCGACCTCCACCTCCTCCACCACCCTCAGTGAAGCCAGGCTCTTG | |
| L5-pabp | AAACGAGCTCGAATTCATCGATGATATGAATGACATTTCTATGAAAGTTTTCAC |  |
| L6-pabp | CAAAACGACTCGCCTGTGAAG |  |
| L3C-rnh1 | CCTTGAAAATACATCTGGCGA | Tagging of Rnh1 |
| L4C-Gly6-rnh1 | TAATTAACCCGGGGATCCGTCGACCTCCACCTCCTCCACCACCCTCAGAAGCTCCTCGCCGAG | |
